# Cue reliability and the evolution of adaptive plasticity in maternal traits

**DOI:** 10.1101/2025.09.06.674614

**Authors:** Henrique Teotónio, Stephen R. Proulx

## Abstract

Natural populations face challenges from multiple environmental factors, and adaptive phenotypic plasticity is expected to evolve only when these factors are strongly correlated across time. In variable environments where developing individuals cannot directly perceive future conditions, natural selection may favor the use of arbitrary but reliable cues by mothers to signal their progeny’s challenging environments, resulting in adaptation that otherwise could not be achieved. We performed evolution experiments with genetically-variable and predominantly selfing populations of the nematode Caenorhabditis elegans in a temporally uncorrelated anoxic selection environment, where during oogenesis and ovulation mothers received blue light pulses that either reliably or unreliably cued impending anoxic conditions in their progeny. After 60 non-overlapping generations of experimental evolution, populations reliably exposed to light evolved both a cue-dependent increase in offspring survival in anoxia following maternal light exposure, and a larger light-independent increase in survival that depended primarily on the maternal oxygen environment. Differentiation between reliable and unreliable populations in their ability to respond to light was most apparent after two generations of anoxia, with evidence of maladaptation in unreliable populations through decreased fecundity and survival in anoxia, although most fitness differentiation between reliable and unreliable populations was observed regardless of light variation, particularly when maternal generations were under normoxia. These results show that when maternal and offspring selection environments are uncorrelated, maternal traits evolve only if maternal environments provide cues predictive of offspring selection. They indicate that cue reliability can determine the evolution of adaptive plasticity in traits with maternal effects.

## Introduction

Whether the evolution of phenotypic plasticity underlies adaptation to temporally variable environments depends on how strong the correlation is between the environmental factor cueing individual development and the environmental factor causing selection (Wade and Kalisz, 1990; Moran, 1992; DeWitt and Scheiner, 2004; Tufto, 2000; Chevin et al., 2010; McNamara et al., 2016). However, in many circumstances, individuals cannot directly detect the relevant environmental state during development and they must instead rely on their parents, typically their mothers, for signals to guide their expression of appropriate trait values in the environment they will experience later in life.

Many empirical examples and theoretical models illustrate how the evolution of traits with maternal effects on offspring can underlie adaptation to temporally variable environments (e.g., Roach and Wulff, 1987; Lande and Kirkpatrick, 1990; Wolf and Brodie, 1998; Mousseau and Fox, 1998; Wolf and Wade, 2009; Marshall and Uller, 2007; McGlothlin and Galloway, 2014; Dey et al., 2016; Proulx and Teotónio, 2017). Maternal effect traits can contribute to adaptation when there is a predictable relationship between the maternal environment and the offspring environment, in particular if the maternal phenotype generates bias in offspring traits toward their optima. More fundamentally, selection creates a positive covariance between the alleles individuals pass on to their progeny and the environment their mothers have experienced (Lande and Price, 1989; Wolf and Wade, 2009, 2016).

For instance, in the nematode *Caenorhabditis elegans*, anoxic conditions early in life lead to embryo mortality when mothers, having accumulated glycerol to withstand osmotic stress, are unable to provision their embryos with enough glycogen for completing development and hatching as free-living larvae (Frazier and Roth, 2009; LaMacchia et al., 2015). Evolution experiments in *C. elegans* conducted by Dey et al. (2016) found that in environments characterized by constant high osmotic stress and temporally alternating anoxia and normoxia, where oxygen level was negatively autocorrelated between maternal and offspring generations, adaptation occurred through the evolution of plasticity in maternal glycogen provisioning. Mothers from evolved populations that developed in normoxia provisioned their embryos with more glycogen than mothers from the ancestral population to ensure progeny survival in anoxia, but did not do so when they developed under anoxia.

If the environmental factor that causes selection is not temporally correlated across generations and there is no additional information available to cue development, mothers cannot signal the impending conditions to their progeny. Adaptation to these variable environments can still occur but would have to rely on an evolutionary response in the variance of trait distributions (Bull, 1987; Seger and Brockmann, 1987; Tufto, 2015; Proulx and Teotónio, 2017); for example if maternal trait expression shows a diversifying “bet-hedging strategy” where individual mothers increase the variance in protection or provisioning among their progeny allowing for a few to survive the challenging conditions at the cost of a reduction in average progeny survival in more benign conditions (Agrawal et al., 1999; Moran et al., 2010; Simons, 2011; Bonduriansky et al., 2016; Joschinski and Bonte, 2020). However, in the *C. elegans* evolution experiments of Dey et al. (2016), populations exposed to temporally uncorrelated anoxia between maternal and offspring generations showed neither evolution of the maternal variance in glycogen provisioning nor evidence of adaptation.

Bet-hedging maternal strategies are expected to have low heritabilities because they increase the environmental component of the trait’s distribution, and in environments with small temporal autocorrelations, selection of maternal trait variance is weak (Crean and Marshall, 2009; Kuijper and Hoyle, 2015; Proulx and Teotónio, 2017). Traits with maternal effects are further known to generate a recursive dependence on their past expression, producing environmental influences that attenuate with generational distance (Lande and Kirkpatrick, 1990; McGlothlin and Galloway, 2014; Kuijper et al., 2014; Wolf and Wade, 2009, 2016). The rate at which maternal effects persist depends on the specific mechanisms of (offspring) development, from the energetic resources provisioned being used during the lifetime of the individual to small RNA regulation, histone post-translational modifications or mitochondrial functions that can endure in some of the more distant descendants (Heard and Martienssen, 2014; Bell and Hellmann, 2019; Zilio et al., 2026). As a consequence, in environments with temporal cross-correlations between environmental factors, or that are temporally autocorrelated over time-scales longer than one generation, selection could also favor the evolution of maternal traits in the lineages that carry over the “memory” of past environments (Jablonka et al., 1995; Day and Bonduriansky, 2011; Furrow and Feldman, 2014; Donelson et al., 2018; Leimar and McNamara, 2015).

The sensitivity of a genotype’s trait expression to environmental variation, i.e., the slope of its reaction norm or “plasticity,” is transiently expected to be under selection when the population is in a novel environment for a few generations, whereas the population’s mean, or the elevation of the reaction norm, is initially expected to change to a lesser extent (Lande, 2009). However, whether the different components of the trait’s genotype-by-environment interaction can be accurately estimated or have independent genetic bases has not been resolved (e.g., Via et al., 1995; Lynch and Walsh, 1998; DeWitt and Scheiner, 2004; Ghalambor et al., 2007; Walsh and Lynch, 2018); and even for traits without maternal effects, how selection determines the evolution of the genotype-by-environment interaction is expected to be complex (Lande, 2009; Michel et al., 2014; Tufto, 2015; Guzella et al., 2018; Draghi, 2023). Nevertheless, theoretical results indicate that if there is no correlation between the maternal and offspring selection environments, adaptive plasticity in maternal traits can only evolve when there is another environmental factor that is itself correlated with the offspring selection environment (Wong and Ackerly, 2005; Kuijper et al., 2014; Chevin and Lande, 2015). Here, we experimentally test this hypothesis by creating situations with an uncorrelated selection environment that had a “reliable cue”. Historically, researchers discuss such cues as environmental states that have no direct effect on fitness, or more technically, that the effect of selection is conditionally independent of the effect of the cue (Pearl, 2009; McNamara et al., 2016; Scheiner, 2018).

We ask how selection in temporally uncorrelated maternal-offspring selection environments may result in adaptation. This could occur with the evolution of a maternal trait detecting an arbitrary environmental cue which in turn triggers the evolution of another maternal trait that increases offspring survival in the selection environment. More generally, adaptation could arise through evolutionary changes in multiple components of the maternal response, including cue-dependent responses as well as other maternal-environment-dependent effects. We experimentally evolved populations of *C. elegans* in an uncorrelated oxygen level time series between maternal and offspring generations. The experiments are similar to those of Dey et al. (2016) but they were designed so that during oogenesis and ovulation, and before reproduction, mothers received one of two light-environment treatments. They either received blue light pulses or no light. In the reliable experimental evolution regime, the light-environment was highly correlated with the offspring oxygen environment, and the blue light pulses were a reliable cue for selection in anoxic conditions (and conversely no light was a reliable cue of normoxia). In the unreliable regime, the maternal light-environment was uncorrelated with the offspring oxygen environment.

*C. elegans* are able to discriminate blue light (Ward et al., 2008; Edwards et al., 2008; Ghosh et al., 2021), avoiding the light source without adverse effects on survival or reproduction when exposed for short periods during oogenesis and ovulation (Proulx et al., 2019). Because individual survival in anoxia depends on maternal glycogen provisioning (Frazier and Roth, 2009; LaMacchia et al., 2015), we assume that during the experiment there is a fixed one-to-one mapping between provisioning and fitness. If after experimental evolution fitness differs between reliably-cued and unreliably-cued populations as a function of light variation, we can also infer that there was evolution of a maternal trait involved in the detection of light prior to provisioning (McNamara et al., 2016; Scheiner, 2018; Nielsen and Papaj, 2022).

Our evolution experiments address several theoretical predictions. Chevin and Lande (2015) found that *de novo* plastic responses of a trait that only signals selection are favored over plastic responses of the trait under selection when the environmental factor cueing development is correlated with the environmental factor causing selection. When explicitly modeling the evolution of maternal traits, Kuijper et al. (2014) found that cross-correlations between environmental factors create variable time-lags of selection, with traits under earlier selection during the lifetime of mothers being more informative about their progeny’s environment. Therefore, we expect that in reliably-cued populations, a maternal light-detection trait will be favored by selection so that mothers can provision their progeny for anoxia when exposed to light before reproduction. Furthermore, by design, there were small autocorrelations of oxygen levels and small cross-correlations between light and oxygen variation at lags longer than maternal–offspring generations. Hence, for adaptation to occur in unreliable populations it must involve either selection for increased maternal variance in provisioning and/or selection of maternal traits that depend on the oxygen environmental states transmitted by ancestors such as grandmothers.

## Materials and Methods

Detailed materials and methods can be found in Supplementary Materials.

### Ancestral population and standard culture conditions

All populations used for experimental evolution were derived from a single ancestral population (GA250). This population originated from the hybridization of 16 inbred founder strains (Teotónio et al., 2012; Teotónio et al., 2017), followed by 140 generations of laboratory domestication (Teotónio et al., 2012; Chelo and Teotónio, 2013; Carvalho et al., 2014) and 50 generations of adaptation to high-salt conditions (305 mM NaCl) from the L1 larval stage through reproduction (Theologidis et al., 2014; Chelo et al., 2019).

During domestication and salt adaptation, populations were maintained at census sizes of *N* = 10^4^ (effective size *N*_*e*_ ≈ 10^3^; Chelo and Teotónio, 2013) on a 4-day life cycle (Teotónio et al., 2012). Populations were cultured on NGM-lite agar plates seeded with *HT115 E. coli*. After 66 h at 20^°^C and 80% relative humidity, adults were bleached (1 M KOH, 5% NaOCl), leaving only embryos. Embryos were incubated for 18 h in M9 buffer without food to obtain synchronized L1 larvae, which were counted and reseeded to initiate the next generation. Generations were therefore non-overlapping.

Although *C. elegans* hermaphrodites can self-fertilize, outcrossing occurs in the presence of males (Stewart and Phillips, 2002). During laboratory domestication, males were maintained at frequencies of 20–40%, corresponding to outcrossing rates of 40–80% (Teotónio et al., 2012; Carvalho et al., 2014). Adaptation to high salt greatly reduced male frequency (Theologidis et al., 2014), but sufficient standing genetic variation remained for evolutionary responses through the sorting of selfing lineages (Noble et al., 2017; Dey et al., 2016). Genome-wide SNP data estimated approximately 30 segregating lineages at intermediate frequencies in the ancestral population (Guzella et al., 2018).

### Oxygen and blue light culture conditions

The first environmental factor manipulated during experimental evolution and the subsequent fitness and fecundity assays (Supplementary Figure S1) was atmospheric oxygen during embryogenesis. After bleaching, embryos were transferred to NGM-lite plates without *E. coli*. Anoxic conditions were imposed from embryogenesis until the L1 stage by incubating plates for 16 h in sealed chambers containing GasPak™ EZ sachets (Becton Dickinson), producing oxygen levels below 1% O_2_, as verified with anaerobic indicator strips. Control plates were maintained under normoxia. Following incubation, live L1 larvae were recovered in M9 buffer, counted, and used to initiate the next generation. Embryonic survival under anoxia depends largely on maternally provisioned glycogen, although embryonic transcription and translation begin before hatching (Bowerman, 1998; Frazier and Roth, 2009; LaMacchia et al., 2015).

The second manipulation was maternal exposure to blue light during oogenesis and ovulation. Beginning 48 h after L1 seeding, adults were exposed for 12 h to 465 nm blue LEDs flashing for 0.5 s every 2 s. The intermittent illumination minimized heat accumulation while providing uniform exposure across plates. Control populations were maintained simultaneously in darkness. Although adult *C. elegans* normally avoid light (Ward et al., 2008; Edwards et al., 2008), individuals in our experimental setup could not escape illumination, ensuring comparable exposure among individuals.

### Experimental evolution design

The cryopreserved ancestral GA250 population was thawed, expanded for two generations, and divided into replicate populations for experimental evolution (Supplementary Figure S2).

The design of the fluctuating oxygen environments has been described previously (Proulx et al., 2019). Briefly, populations experienced 60 generations comprising 30 generations in normoxia and 30 in anoxia. Three environmental sequences were used that differed in the temporal autocorrelation of anoxic conditions between grandmaternal and grand-offspring generations, with temporal autocorrelations at higher lags being negligible (Seq19, Seq31, Seq32; Supplementary Figure S2, Supplementary Methods).

Within each oxygen sequence, maternal blue-light exposure either reliably or unreliably predicted offspring anoxia (Supplementary Figure S2). In the reliable regime, maternal light perfectly predicted anoxia in the offspring generation, whereas in the unreliable regime the probability was 0.5. The two regimes therefore differed only in the correlation between maternal light exposure and offspring anoxia. Both regimes experienced identical numbers of anoxic generations and maternal light exposures over the course of the experiment. Cross-correlations at higher lags were weak.

For each combination of oxygen sequence and cue reliability, three replicate populations were maintained under standard laboratory conditions (305 mM NaCl; census size *N* = 10^4^) for 60 generations. This yielded 18 independently evolved populations: nine evolved under reliable environmental cues and nine under unreliable cues. Population identities are listed in the detailed Supplementary Methods.

### Fitness and fecundity assays

Fitness was measured as the discrete-time population growth multiplier, *R*_0_, under anoxic offspring conditions (see next section). Specifically, *R*_0_ was calculated as the number of live L1 larvae surviving anoxia, after grandmaternal and maternal oxygen treatments and maternal light treatment (Supplementary Figure S1), divided by the fixed number of L1 larvae seeded in the maternal generation.

Fitness assays were performed after experimental evolution using common-garden conditions. Eighteen assay blocks were conducted, each including the ancestral population (GA250) and one evolved population. Within each block, all eight combinations of grandmaternal oxygen, maternal oxygen, and maternal light treatments were assayed. Each population was thawed from cryopreserved stocks, expanded for two generations under standard laboratory conditions, and then exposed to the appropriate environmental treatments beginning in the grandmaternal generation. Three technical replicates were performed for every environmental combination. The total number of surviving offspring L1 larvae after anoxia was estimated from subsamples and divided by the number of seeded larvae in the maternal generation to obtain *R*_0_. The complete dataset comprised 845 observations.

Fecundity assays were performed separately in six assay blocks using the ancestral population together with the six evolved populations from Seq31. Fecundity was measured as the number of embryos produced after the grandmaternal and maternal environmental treatments, but before offspring exposure to anoxia (Supplementary Figure S1), divided by the number of seeded maternal L1 larvae. All other aspects of the design, including common-garden culture, technical replication, and assay procedures, matched those used for the fitness assays. The fecundity dataset comprised 285 observations.

For both assays, all experimental manipulations were performed by the same experimenter using randomized sample identities to minimize handling bias. Population samples experienced five generations of common culture before measurement, minimizing uncontrolled environmental carry-over effects. Because some adults and offspring were unavoidably lost during bleaching and handling, both *R*_0_ and fecundity should be regarded as conservative (lower-bound) estimates of their true values.

### Fitness analysis of the ancestral population

The discrete-time population growth multiplier, *R*_0_, was used as a proxy for fitness. Analyses were performed on the Malthusian scale, *w* = ln(*R*_0_), which is approximately equivalent to the per-generation population growth rate under our experimental design (Otto and Day, 2007). Results are presented on the natural *R*_0_ scale.

We first modeled the ancestral population to estimate baseline fitness across the eight assay environments defined by the combinations of grandmaternal oxygen, maternal oxygen, and maternal light treatments (Figure 1). The model included environmental-state effects and a random effect of assay block:

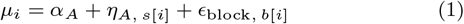

where *α*_*A*_ is the overall ancestral mean fitness, *η*_*A,s*_ is the effect of assay state *s*, and *ϵ*_block_ accounts for variation among assay blocks. State effects were estimated independently (equivalent to fitting all interactions among grandmaternal oxygen, maternal oxygen, and maternal light environments) and constrained to sum to zero to ensure that they represent deviations from the group means.

**Figure 1.**
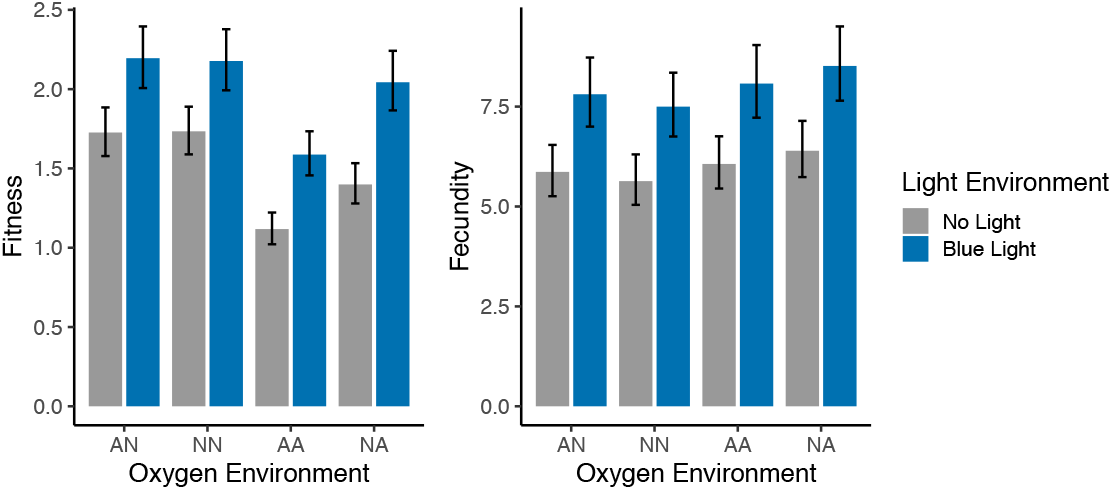
Fitness and fecundity of the ancestral population. Left. Fitness under anoxic conditions was measured as the per capita population growth rate, *R*_0_, following grandmaternal and maternal exposure to normoxia (N) or anoxia (A), and maternal exposure to either blue light pulses (blue) or no light (gray) during oogenesis and ovulation (Materials and Methods, Supplementary Figure S1), with two-letter labels denoting grandmaternal and maternal oxygen environments, respectively. Measurements were made in 18 independent assay blocks, each with 3 technical replicates (Materials and Methods). Right. Fecundity of the ancestral population, prior to offspring exposure to anoxia (Supplementary Figure S1). Measurements were made in 8 independent assay blocks, each with 3 technical replicates (Materials and Methods). For both panels, the bars show posterior medians of the estimates after back-transforming to the original measurement scale, with 95% probable interval (PI) for the ancestral population (equation (1) for fitness, and equation (3) for fecundity; see Materials and Methods).

### Fitness analysis of reliable and unreliable populations

We next modeled evolutionary changes in fitness relative to the ancestral population. The model partitions evolutionary responses into changes in mean fitness and changes in environmental state-specific fitness, allowing us to distinguish overall evolutionary divergence from changes in fitness plasticity. Reliable and unreliable populations were modeled as deviations from the ancestral population, while accounting for assay block, environmental sequence, and replicate population effects:

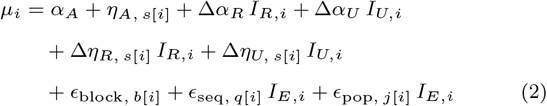

Here, Δ*α*_*R*_ and Δ*α*_*U*_ represent evolutionary changes in mean fitness relative to the ancestral population, whereas Δ*η*_*R*_ and Δ*η*_*U*_ describe changes in fitness specific to each assay environment (Figure 2). These state-specific terms were estimated independently, equivalent to fitting all interactions among grandmaternal oxygen, maternal oxygen, maternal light, and evolutionary regime.

**Figure 2.**
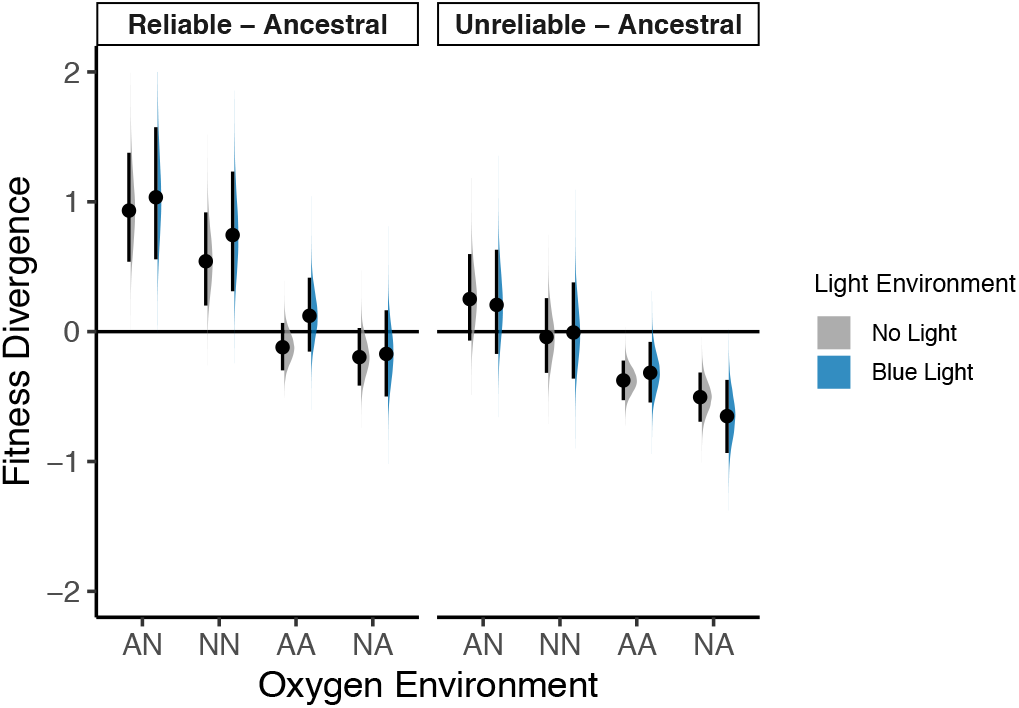
Fitness divergence from ancestral. Fitness differences from the ancestral population of reliable and unreliable populations after 60 generations of evolution under temporally uncorrelated oxygen conditions (Supplementary Figure S2). Fitness was measured in the offspring generation under anoxia, following grandmaternal and maternal exposure to normoxia (N) or anoxia (A), and maternal exposure to either blue light pulses (blue) or no light (gray) during oogenesis and ovulation (Supplementary Figure S1), with two-letter labels denoting grandmaternal and maternal oxygen environments, respectively. Posterior estimates are shown as median differences from the corresponding ancestral environmental states, with median and 95% PIs estimates (dot and interval). Half-eye plots show the posterior distributions for each comparison, calculated as the difference between evolved and ancestral posterior fitted values on the natural scale. On the log-fitness scale of the Bayesian model, evolved populations differ from the ancestral population only through the sum of the Δ*α* and Δ*η* terms in equation (2) (see Materials and Methods). Each oxygen and light treatment combination includes 9 populations plus the ancestral. Posterior distributions are based on 18 independent assay blocks, each with 3 technical replicates. See also Supplementary Figure S3 for results including the ancestral population, and Supplementary Figure S4 for the variance terms of the model.

Assay block, environmental sequence, and replicate population were included as hierarchical random effects. Population effects were modeled separately for the reliable and unreliable regimes but shared a common variance parameter, allowing direct interpretation of the evolutionary effects. As in the ancestral model, state effects and random effects were constrained to sum to zero.

Weakly informative regularizing priors were used throughout, and models were fit by Hamiltonian Monte Carlo (details in Supplementary Materials).

### Fecundity analysis

Fecundity was analyzed using a model analogous to the fitness model, with two modifications. First, because fecundity was measured only for populations evolved under Seq31, the sequence-level random effect was omitted. Second, unlike fitness, which was analyzed as the log-transformed response *w* = log(*R*_0_), fecundity was analyzed on its natural scale using a Gamma likelihood with a log link. This formulation accommodates positive-valued responses whose variance may increase with the mean. A log-normal model yielded nearly identical parameter estimates and conclusions.

Preliminary analyses found no evidence for interactions between oxygen and light environments, so grandmaternal and maternal oxygen effects and maternal light effects were modeled additively. Light treatment was contrast coded (−1 = no light, +1 = light).

The model specification was

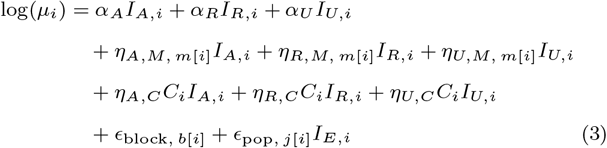

Here, *α* parameters represent regime-specific mean fecundity on the log scale, while the *η* parameters describe the effects of oxygen and light environments. Assay block and replicate population were included as hierarchical random effects, with replicate populations in the reliable and unreliable regimes sharing a common variance parameter. Sum-to-zero constraints were imposed where appropriate for parameter identifiability.

Priors followed the same weakly informative scheme as for fitness and are specified in Supplementary Materials. The fitted model estimates regime-specific mean fecundity and environmental responses while accounting for variation among assay blocks and replicate populations (Figure 4).

**Figure 3.**
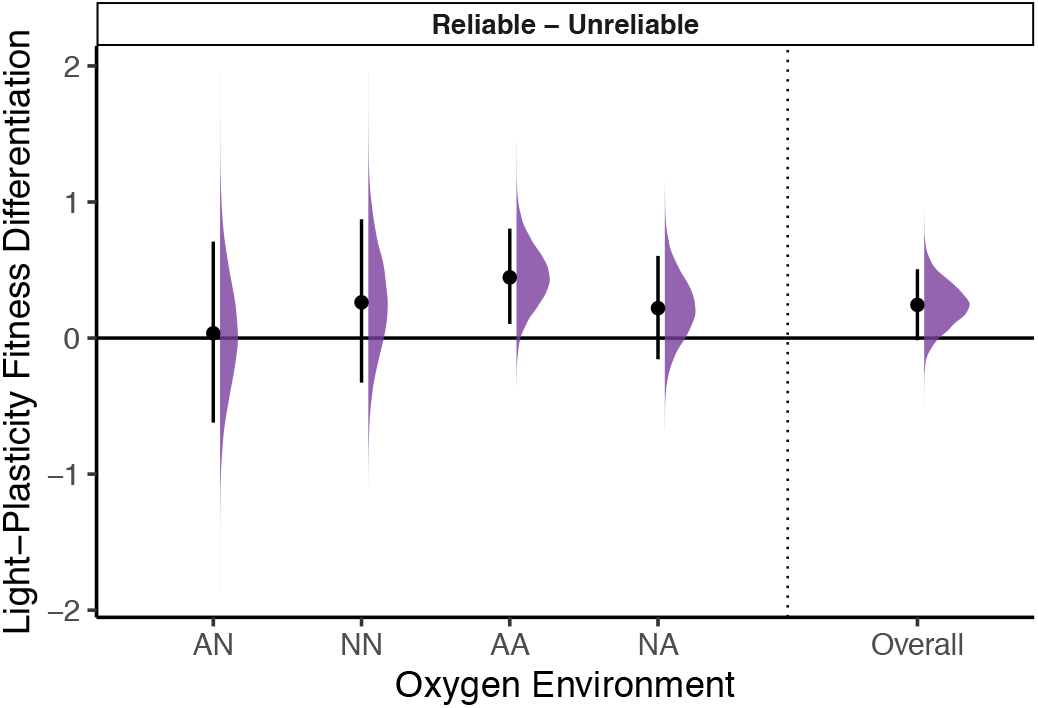
Evolutionary differentiation of fitness plasticity to light variation. Evolutionary differentiation in light-plasticity, measured as the Reliable-Unreliable difference in the fitness effect of light presence versus absence. Half-eye plots show posterior distributions of the Reliable-Unreliable difference in light-plasticity, calculated for each oxygen environment as the difference between the regime-specific natural-scale light responses, [exp(*µ*_*R,m*,Light_) − exp(*µ*_*R,m*,No Light_)] − [exp(*µ*_*U,m*,Light_) − exp(*µ*_*U,m*,No Light_)], where *µ*_*T,s*_ = *α*_*A*_ + *η*_*A,s*_ + Δ*α*_*T*_ + Δ*η*_*T,s*_ (equations (4) and (5), see Materials and Methods) for each oxygen environment. The overall contrast is the average across oxygen environments. The plots show the median and 95% PIs posterior estimates (dot and interval). Positive values indicate increased differentiation in light-plasticity of reliable populations as compared with unreliable populations. “N” indicates normoxia and “A” indicates anoxia, with two-letter labels denoting grandmaternal and maternal oxygen environments, respectively. We also present the average differentiation in light-plasticity across oxygen environments labelled as “Overall”. See Supplementary Figure S5 for the posterior distributions of light-plasticity for each regime, including the ancestral population, and Supplementary Figure S6 for the evolutionary divergence of reliable and unreliable populations relative to the ancestral.

**Figure 4.**
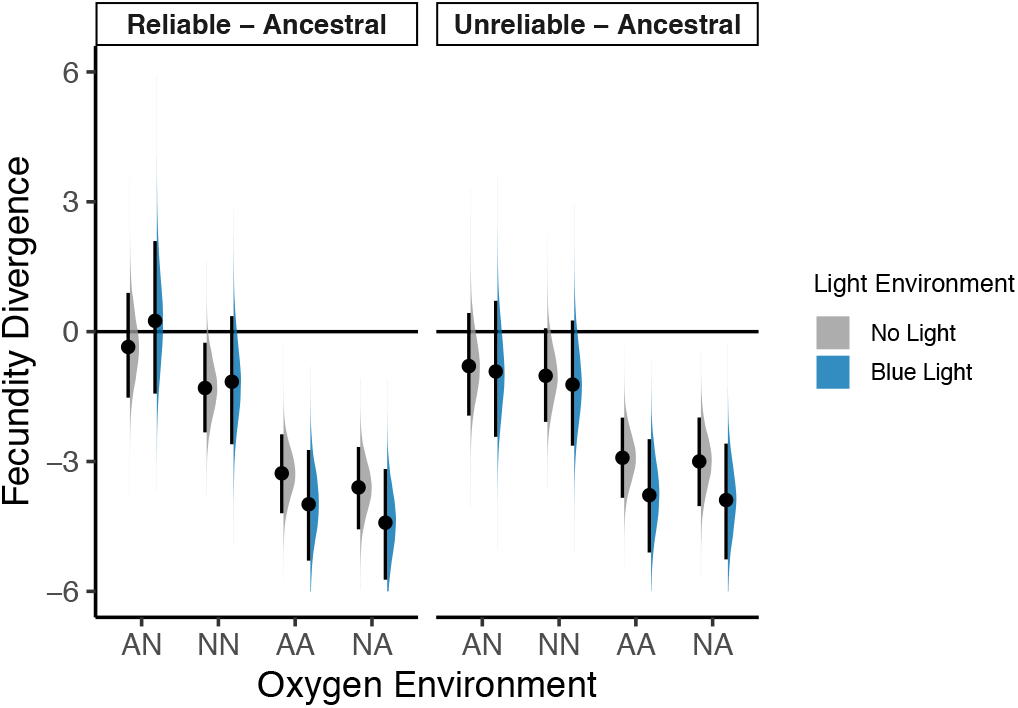
Fecundity divergence from ancestral. Fecundity was measured as the number of embryos per individual mother, following grandmaternal and maternal exposure to normoxia (N) or anoxia (A), and maternal exposure to either blue light pulses (blue) or no light (gray) during oogenesis and ovulation (Supplementary Figure S1), with two-letter labels denoting grandmaternal and maternal oxygen environments, respectively. Posterior estimates are shown as median differences from the corresponding ancestral environmental states, with median and 95% PIs estimates (dot and interval). Half-eye plots show the posterior distributions for each comparison (the *η*_*R*_ and *η*_*U*_ terms in equation (3)). Each oxygen and light treatment combination includes 3 replicate populations plus the ancestral. Posterior distributions are based on 6 independent assay blocks, each with 3 technical replicates. See also Supplementary Figure S7 for results including the ancestral population, Supplementary Figure S8 for the variance terms of the model, and Supplementary Figure S9 for average fecundity regime divergence.

### Contrasting evolutionary responses

Evolutionary changes in the mean elevations of the fitness reaction norms are captured by the Δ*α*_*R*_ and Δ*α*_*U*_ terms in equation (2), whereas environmental state-specific evolutionary changes are described by the Δ*η*_*R*_ and Δ*η*_*U*_ terms (Figure 2). To quantify evolutionary changes in the light-dependent component of fitness plasticity, we contrasted the Δ*η* terms on the natural (*R*_0_) scale. Because fitness and fecundity are interpreted and presented throughout on their natural scales, these contrasts measure absolute changes in offspring survival or embryo production. Note that contrasts on the log scale would describe multiplicative changes, i.e. ratios.

For each posterior draw, we first computed the expected fitness under each environmental state using the fitted linear predictor (equation (2)):

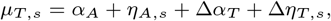

where *T* denotes the Reliable or Unreliable evolutionary regime. Block, sequence, and population random effects were set to zero (their mean values), such that Δ*α*_*T*_ + Δ*η*_*T,s*_ represents the evolutionary divergence from the ancestral population for environmental state *s*.

The fitness response to maternal light exposure was then calculated for each oxygen environment as

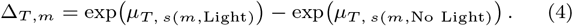

Evolutionary differentiation in light plasticity between the reliable and unreliable regimes was quantified as

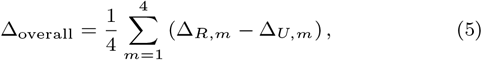

computed for each posterior draw to obtain its posterior distribution (Figure 3). This quantity is a difference of differences on the natural scale; a value of zero indicates no evolutionary differentiation in light plasticity between the two experimental evolution regimes.

An analogous contrast was computed for fecundity, in which oxygen and light effects were modeled additively so that light plasticity could be evaluated at the average oxygen effect; its full derivation is given in Supplementary Materials.

### Software

All fitness and fecundity models were coded using the rethinking package (McElreath, 2020) and run using R (4.3.1) and Stan (2.32.2) (R Core Team, 2021; Stan Development Team, 2020). ChatGPT (GPT-5.3 model) was used for R figure code generation; prompts were structured as natural-language queries specifying the task, and all outputs were reviewed.

## Results

### Environmental effects in the ancestral population

The ancestral population used for experimental evolution was previously adapted to standard laboratory conditions, including normoxia during embryogenesis and hatching at the first larval stage (L1), high salt concentration (305 mM NaCl) in the growth medium from L1 to reproduction, and absence of blue light pulses throughout the life cycle (Teotónio et al., 2012; Theologidis et al., 2014). This ancestral population was derived from a hybrid of 16 inbred founders maintained for almost 200 generations at high population sizes (*N* = 10^4^, *N*_*e*_ = 10^3^) under non-overlapping generations and high outcrossing rates when in low salt conditions, ensuring that standing genetic variation was available for selection responses before the onset of experimental evolution (Chelo and Teotónio, 2013; Noble et al., 2017; Guzella et al., 2018).

The ancestral population was assayed for the *per capita* population growth multiplier in anoxia as a proxy for survival under anoxic conditions, and which we call fitness (Supplementary Figure S1). Assays followed grandmaternal and maternal exposure to variable oxygen levels during development and larval hatching, and maternal exposure to blue light pulses during oogenesis and ovulation prior to reproduction and offspring exposure to anoxia. There were thus 8 environmental states to be compared (see Materials and Methods).

When we compare among oxygen and light treatments, we find that exposure to light increases fitness relative to the dark treatment (Figure 1). This direct effect of light was the largest environmental effect we detected in the ancestral population, exceeding the effects of the grandmaternal and maternal oxygen environments. The maternal and grandmaternal oxygen environments had small effects on fitness, except under dark conditions and when two generations of anoxic conditions were experienced or when mothers developed under anoxia, both of which resulted in reduced fitness.

The ancestral population was also assayed for fecundity. Fecundity was measured as the number of embryos after grandmaternal oxygen and maternal oxygen and light treatments, and thus before offspring exposure to anoxia as in the fitness assays (Supplementary Figure S1). We find similar results as for fitness, in that light exposure increases nematode fecundity (Figure 1). However, and unlike fitness, there are no obvious interactions between light and oxygen environments in determining fecundity.

### Evolutionary divergence from ancestral

Experimental evolution proceeded for 60 generations under uncorrelated oxygen conditions between maternal and offspring generations (autocorrelation of oxygen level time series at lag 1: *ρ*_1_ = −0.065), with three variable grandmaternal to grand-offspring correlations (−0.311 *< ρ*_2_ *<* 0.247). Blue light pulses from which individuals could not escape were given during oogenesis and ovulation (or their absence), serving as either a reliable or unreliable cue of anoxia (or normoxia) in the following generation (Supplementary Figure S2). For each combination of light and oxygen regimes, three replicate populations were derived from the lab-domesticated ancestral population, yielding 9 reliable and 9 unreliable populations with respect to light regime (Materials and Methods). After 60 generations, reliable and unreliable populations were compared in common garden assays to the ancestral population for fitness in anoxia (Supplementary Figure S1).

Reliable populations showed increased fitness when compared to the ancestral population only when mothers developed under normoxia, irrespective of light treatment (Figure 2, Supplementary Figure S3). In contrast, unreliable populations showed evolution of reduced fitness, particularly when assayed under maternal anoxia but also irrespective of light treatment. While the different environmental sequences of oxygen states each population faced during experimental evolution contributed to the variance in fitness, their consequences were relatively smaller than those from other sources (Supplementary Figure S4). Replicate population effects, block effects and intrinsic noise, including technical assay error, were of comparable magnitude.

### Evolution of fitness plasticity to light variation

We next examined the evolution of the fitness difference in the sensitivity to the light assay treatment, light or no light, which we term light-plasticity for fitness (see Materials and Methods). Regardless of oxygen variation, we find that reliable populations had small increases in light-plasticity when compared to the ancestral while unreliable populations had small decreases (Supplementary Figure S5). Similar results are found when reliable and unreliable populations are contrasted to the ancestral population in all oxygen assay environments, with noticeable large variance estimates (Supplementary Figure S6).

Despite little evolutionary divergence in light-plasticity from the ancestral population in each oxygen environment, there is fitness differentiation between the reliable and unreliable populations (Figure 3). This is most evident when populations were assayed under grandmaternal and maternal anoxia, where reliable populations evolved to be more plastic in fitness with regards to light variation than unreliable populations.

### Evolution of fecundity

We also measured fecundity in a subset of three replicate populations from each of the reliable and the unreliable regimes, those where the temporal autocorrelation in oxygen levels between grandmaternal and grand-offspring generations was positive during experimental evolution (*ρ*_2_ = 0.247, Materials and Methods).

We find that reliable and unreliable populations have diverged from the ancestral population by decreasing fecundity, particularly when mothers developed in anoxia (Figure 4, Supplementary Figures S7 and S8). On average reliable and unreliable populations diverged similarly from the ancestral by reducing fecundity (Supplementary Figure S9). Differentiation between reliable and unreliable populations in their plasticity for fecundity under the light environment was also minimal, independently of oxygen environments (Supplementary Figures S10).

### Adaptation to uncorrelated oxygen environments

Under fluctuating environments, the hallmark of adaptation is an increase in the geometric mean fitness across the sequence of oxygen and light environmental states that a population experienced during its own evolutionary history (Cohen, 1966; Gillespie, 1974; Proulx and Day, 2001; Lande, 2007; Saether and Engen, 2015). To measure adaptation, we averaged the log of the fitness estimates across all combinations of grandmaternal and maternal environments, taking into account that both environmental factors (light and oxygen) were equally frequent in the evolution experiments and that there were small autocorrelations between maternal and offspring oxygen environments (Supplementary Figure S2, Materials and Methods). For the reliable populations, adaptation was calculated only when blue light pulses were given to mothers and their progeny experienced anoxia. For the unreliable populations, we calculated adaptation under the conditions that they were exposed to which included light half of the time.

We find that reliable populations adapted to their uncorrelated maternal-offspring oxygen environments, while unreliable populations showed maladaptation (Figure 5, Supplementary Figure S11).

**Figure 5.**
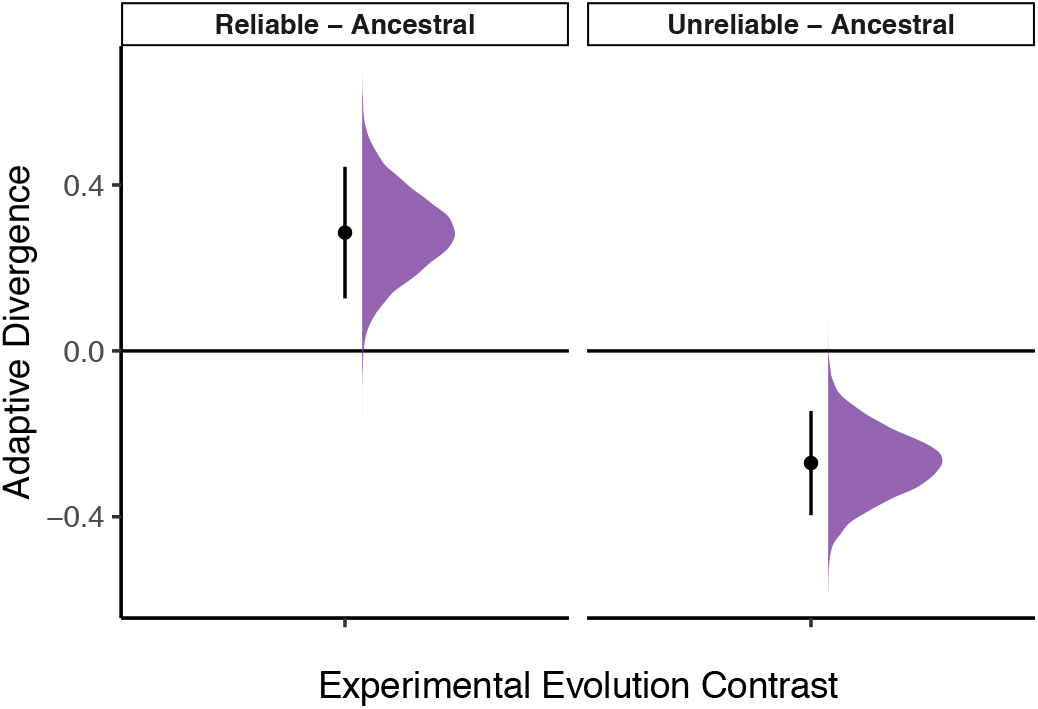
Adaptive divergence from ancestral. To measure adaptation, we averaged the log of posterior fitness estimates over all combinations of grandmaternal and maternal environments the populations experienced during experimental evolution. This corresponds to a calculation of the geometric mean fitness on the measured scale. Posterior estimates are shown as median differences from the ancestral population, with median and 95% PIs estimates (dot and interval). Half-eye plots show the posterior distributions for each comparison. See Supplementary Figure S11 for the posterior distributions of each experimental regime including the ancestral population.

## Discussion

If there is no correlation between the maternal and offspring selection environments, adaptive plasticity in maternal traits can only evolve when another environmental factor is itself correlated with the offspring selection environment, and our evolution experiments support this theoretical prediction (see Introduction). In the reliable regime, the light environment was strongly correlated with the offspring oxygen environment, such that blue light pulses during maternal oogenesis and ovulation reliably cued anoxic selection in offspring generations, whereas in the unreliable regime, the maternal light environment was uncorrelated with offspring oxygen conditions. Populations evolving with reliable cues adapted to the fluctuating oxygen environment, while those evolving under unreliable cues showed maladaptation (Figure 5).

The cue-reliability regimes experienced identical oxygen sequences and identical numbers of light-exposed generations, differing only in the correlation between them. Selection on traits without maternal effects, including those involved in light detection and anoxia survival, must therefore have been equivalent between regimes, and the adaptive divergence we observe is attributable to maternal traits. This inference further requires that the differences we measure among the ancestral, reliable, and unreliable populations are themselves evolved (genetic) rather than arising from uncontrolled carry-over from the specific conditions each population sample last experienced. Population samples, including the ancestral, were thawed from cryopreserved stocks and cultured under identical conditions for five generations, and shared the same randomized handling and oxygen or light treatments (Materials and Methods). Any systematic differences in fitness or fecundity should therefore reflect evolved genetic differences rather than short-term physiological state.

The observed adaptive responses must be qualified, however, because survival in normoxia was not measured, and a trade-off with anoxia survival cannot be excluded. We therefore cannot rule out that reliable populations evolved reduced fitness in normoxia, or that the response to uncorrelated selection environments with an unreliable light cue involved increasing fitness under normoxic conditions with reduced fitness in anoxia. Both scenarios seem unlikely because the ancestral population adapted to normoxia for nearly 200 generations prior to experimental evolution (Chelo and Teotónio, 2013; Theologidis et al., 2014), and because, at least in the reliable populations, dark conditions cued normoxia, providing little opportunity for the evolution of reduced survival under normoxic conditions. In line with this, reliable populations whose mothers developed in normoxia and faced dark conditions before reproduction showed increased progeny survival in anoxia by almost as much as when mothers were exposed to light (Figure 2). By design, such correlated evolutionary responses in reliable populations could only have occurred because of anoxic selection in offspring generations (Lande and Price, 1989; Wolf, 2000; Wolf and Wade, 2016).

Survival in anoxia is known to depend almost exclusively on maternal glycogen provisioning (Frazier and Roth, 2009; LaMacchia et al., 2015), and so the fitness gains observed in reliable populations likely reflect increased provisioning, as previously shown under negatively alternating normoxia–anoxia experimental evolution regimes (Dey et al., 2016). In the reliable populations, provisioning depended on maternal detection of blue light pulses before reproduction, indicating the evolutionary divergence of genotype-by-environment interactions in maternal traits involved in light detection and provisioning. It is tempting to read this divergence as evolution of the slope of the maternal reaction norm to light alone, following the conventional decomposition of reaction norms into elevations and slopes (e.g., Lande, 2009; Chevin et al., 2010; Tufto, 2015). The components of the genotype-by-environment interaction are, however, neither necessarily genetically-independent nor tunable by selection (Via et al., 1995; Lynch and Walsh, 1998; DeWitt and Scheiner, 2004; Walsh and Lynch, 2018), and in our own models we required sum-to-zero constraints on elevations and hierarchical priors on elevations and slopes to ensure identifiability (Pearl, 2009; Gelman et al., 2013). Our results can nonetheless be described as the evolution of “multivariate plasticity” in maternal traits, because evolutionary responses depended jointly on light and oxygen environmental variation, cf. Nielsen and Papaj (2022).

Reliable populations differed from unreliable populations in two respects, a large increase in the light-independent maternal effect and a modest increase in the response to the light-cue. Maternal light perception and resource provisioning are likely to be under pleiotropic genetic control due to physiological interconnectedness, so evolutionary changes affecting one component are expected to influence others. We speculate that this pattern could arise if maternal responses include physiological changes that persist across generations. Under the reliable regime, where maternal light consistently predicted offspring anoxia, selection could both maintain these inherited maternal effects and improve their overall performance, leading to increases in both baseline offspring fitness and responsiveness to the light cue. In contrast, under the unreliable regime, these inherited effects would more often be mismatched to offspring environments, favoring their reduction.

The fecundity results support the interpretation that the increase in overall fitness reflects improved offspring survival rather than increased reproduction. Fecundity did not show evolved increases in the environmental states where reliable populations showed higher fitness, and reliable populations instead evolved reduced fecundity when mothers had developed in anoxia, independently of light variation (Figure 4). A similar pattern evolved under negatively alternating maternal–offspring normoxia– anoxia environments, where increased glycogen provisioning by normoxia-developed mothers was correlated with reduced fecundity in anoxia-developed mothers (Dey et al., 2016). The corresponding dependence of fitness and fecundity on maternal oxygen state is therefore consistent with a trade-off between progeny number and investment in offspring survival under anoxia. In *C. elegans*, reduced progeny production can also enhance adult survival under oxygen deprivation (Mendenhall et al., 2009). Although we did not measure the underlying physiological mechanism, glycogen allocation to embryos at the expense of oogenesis and ovulation provides one possible explanation for the pattern. Together, the fitness and fecundity responses support the conclusion that the evolutionary changes involve maternal life-history traits (Proulx and Teotónio, 2017; Kuijper and Johnstone, 2021).

Unreliable populations showed maladaptation, with reduced fecundity and reduced fitness (presumably also reduced survival) relative to the ancestral population (Figures 2 and 4). Contrary to theoretical expectations, we found no evidence that unreliable populations adapted through increased variance in maternal provisioning or through maternal effects persisting across multiple generations (see Introduction). This outcome is not surprising: despite the potential for standing genetic variation in traits that carry over the “memory” of past environments —which are known to vary genetically in *C. elegans* (Frézal et al., 2018; Frejacques et al., 2026)—, and although more research on this is needed, there was likely little possibility for such traits to be selected, given the weak oxygen autocorrelations and weak cross-correlations between light and oxygen at lags beyond the maternal-offspring generations (Proulx and Teotónio, 2017; Proulx et al., 2019). Unreliable populations were maladapted relative to the ancestral population, but we estimated their geometric mean fitness as being above 1, which explains why they could persist for 60 generations despite reduced performance under anoxic conditions (Supplementary Figure S11).

The cue-dependent component of the evolutionary responses in fitness and fecundity raises the question of how physiological changes in the germline of grandmothers may have influenced the evolution of light detection in their adult progeny. Light detection in *C. elegans* occurs via ciliated head neurons and unique photoreceptor proteins (Edwards et al., 2008; Ward et al., 2008; Liu et al., 2010). Neuronal gene expression may depend on anoxia exposure during development (Zhang et al., 2020), and the evolution of several neuroendocrine mechanisms could have mediated the evolution of the light signal transduction to the germline (Hubbard et al., 2013; Burton et al., 2017; Das et al., 2020; Zhou and Liu, 2026). Notably, the ancestor population had not previously been exposed to light, yet exhibited increased fecundity and fitness upon exposure (Figure 1), suggesting a hormetic response in which low doses of light confer physiological benefits, whereas higher doses are detrimental, consistent with the aversive response of *C. elegans* to light (Edwards et al., 2008; Ward et al., 2008). Strikingly, our results parallel a recent study showing that light exposure can enhance thermotolerance in individuals and in their offspring, even when the latter are kept in the dark (Zhou and Liu, 2026).

In conclusion, the evolution experiments presented here show that adaptive plasticity in maternal traits evolves when environmental conditions experienced by maternal generations provide reliable cues about the selective environments their offspring are likely to encounter. Although natural populations are continuously exposed to multiple environmental factors, how they adapt to changing conditions remains poorly understood. It is unclear, for instance, whether organisms living in seasonal environments use variation in precipitation or temperature as cues to adjust resource allocation and maximize reproductive success, or whether these same factors also act directly as selection agents. Our findings indicate that natural populations could have evolved to respond to cues from one dimension of their environment in order to adaptively predict changes in another, particularly by the evolution of traits with maternal effects.

## Data and code availability

Data, contrast results, and code for analysis and figure generation are available in our GitHub repository (https://github.com/ExpEvolWormLab/Evolution-Blue-Light-Cue).

## Author Contributions

HT: Conceptualization, Resources, Data curation, Supervision, Funding acquisition, Investigation, Writing, Project administration; SRP: Conceptualization, Software, Formal analysis, Validation, Investigation, Funding acquisition, Writing.

## Acknowledgments

We are indebted to Snigdhadip Dey, a talented biologist who conducted the evolution experiments and performed the fitness and fecundity assays, and who passed away during the course of this work. We also thank H. Gendrot, V. Pereira, S. Santos, A. Larue, S. Hotte and D. Tran for their help with nematode handling, C. Braendle and F. Mallard for discussion, and manuscript referees for their helpful comments.

## References

Agrawal, A. A., Laforsch, C., & Tollrian, R. Transgenerational induction of defences in animals and plants. Nature, 401(6748): 60–63, 1999. 10.1038/43425.

Bell, A. M. & Hellmann, J. K. An integrative framework for understanding the mechanisms and multigenerational consequences of transgenerational plasticity. Annu Rev Ecol Evol Syst, 50:97–118, 2019. 10.1146/annurev-ecolsys-110218-024613.

Bonduriansky, R., Runagall-McNaull, A., Crean, A. J., & Lee, K. P. The nutritional geometry of parental effects: maternal and paternal macronutrient consumption and offspring phenotype in a neriid fly. Functional Ecology, 30(10): 1675–1686, 2016. 10.1111/1365-2435.12643.

Bowerman, B. Maternal control of pattern formation in early caenorhabditis elegans embryos. Curr Top Dev Biol., 39:73–117., 1998. 10.1016/S0070-2153(08)60453-6.

Bull, J. Evolution of phenotypic variance. Evolution, 41:303–315, 1987. 10.1111/j.1558-5646.1987.tb05799.x.

Burton, N. O., Furuta, T., Webster, A. K., Kaplan, R. E., Baugh, L. R., Arur, S., & Horvitz, H. R. Insulin-like signalling to the maternal germline controls progeny response to osmotic stress. Nat Cell Biol, 19(3):252–257, 2017. 10.1038/ncb3470.

Carvalho, S., Phillips, P., & Teotónio, H. Hermaphrodite life history and the maintenance of partial selfing in experimental populations of caenorhabditis elegans. BMC Evol Biol, 14:117, 2014. 10.1186/1471-2148-14-117.

Chelo, I. M. & Teotónio, H. The opportunity for balancing selection in experimental populations of caenorhabditis elegans. Evolution, 67(1):142–56, 2013. 10.1111/j.1558-5646.2012.01744.x.

Chelo, I. M., Afonso, B., Carvalho, S., Theologidis, I., Goy, C., Pino-Querido, A., Proulx, S., & Teotónio, H. Partial selfing can reduce genetic loads while maintaining diversity during experimental evolution. G3 (Bethesda), 9:2811–2821, 2019. 10.1534/g3.119.400239.

Chevin, L. M. & Lande, R. Evolution of environmental cues for phenotypic plasticity. Evolution, 69(10):2767–75, 2015. 10.1111/evo.12755.

Chevin, L. M., Lande, R., & Mace, G. M. Adaptation, plasticity, and extinction in a changing environment: towards a predictive theory. PLoS Biol, 8(4):e1000357, 2010. 10.1371/journal.pbio.1000357.

Cohen, D. Optimizing reproduction in a randomly varying environment. Journal of Theoretical Biology, 12:119–129, 1966.

Crean, A. J. & Marshall, D. J. Coping with environmental uncertainty: dynamic bet hedging as a maternal effect. Philos Trans R Soc Lond B Biol Sci, 364(1520):1087–96, 2009. 10.1098/rstb.2008.0237.

Das, S., Ooi, F. K., Cruz Corchado, J., Fuller, L. C., Weiner, J. A., & Prahlad, V. Serotonin signaling by maternal neurons upon stress ensures progeny survival. eLife, 9:e55246, 2020. 10.7554/eLife.55246.

Day, T. & Bonduriansky, R. A unified approach to the evolutionary consequences of genetic and nongenetic inheritance. Am Nat, 178(2):E18–36, 2011. 10.1086/660911.

DeWitt, T. & Scheiner, S. M. Phenotypic Plasticity: Functional and Conceptual Approaches. Oxford University Press, Oxford, 2004.

Dey, S., Proulx, S., & Teotónio, H. Adaptation to temporally fluctuating environments by the evolution of maternal effects. PLoS Biol, 14(2):e1002388, 2016. 10.1371/journal.pbio.1002388.

Donelson, J. M., Salinas, S., Munday, P. L., & Shama, L. N. S. Transgenerational plasticity and climate change experiments: Where do we go from here? Glob Chang Biol, 24(1):13–34, 2018. 10.1111/gcb.13903.

Draghi, J. A. Bet-hedging via dispersal aids the evolution of plastic responses to unreliable cues. J Evol Biol, 36(6):893–905, 2023. 10.1111/jeb.14182.

Edwards, S. L., Charlie, N. K., Milfort, M. C., Brown, B. S., Gravlin, C. N., Knecht, J. E., & Miller, K. G. A novel molecular solution for ultraviolet light detection in caenorhabditis elegans. PLoS Biol, 6(8):e198, 2008. 10.1371/journal.pbio.0060198.

Frazier, H. N. r. & Roth, M. B. Adaptive sugar provisioning controls survival of c. elegans embryos in adverse environments. Curr Biol, 19(10):859–63, 2009. 10.1016/j.cub.2009.03.066.

Frejacques, F., Saglio, M., Aljohani, M., Chervova, A., Bourdon, L., Pelletier, K., Cecere, G., Froekjaer-Jensen, C., Frézal, L., & Félix, M.-A. Intraspecific variation in the duration of epigenetic inheritance in wild isolates of ¡em¿c. elegans¡/em¿. Current Biology, 36(7):1675–1690.e7, 2026. 10.1016/j.cub.2026.02.040.

Frézal, L., Demoinet, E., Braendle, C., Miska, E., & Félix, M. A. Natural genetic variation in a multigenerational phenotype in c. elegans. Curr Biol, 28(16):2588–2596 e8, 2018. 10.1016/j.cub.2018.05.091.

Furrow, R. E. & Feldman, M. W. Genetic variation and the evolution of epigenetic regulation. Evolution, 68(3):673–83, 2014. 10.1111/evo.12225.

Gelman, A., Carlin, J. B., Stern, H. S., Dunson, D. B., Vehtari, A., & Rubin, D. B. Bayesian Data Analysis. CRC Press, Boca Raton, FL, 3 edition, 2013.

Ghalambor, C. K., McKay, J. K., Carroll, S. P., & Reznick, D. N. Adaptive versus non-adaptive phenotypic plasticity and the potential for contemporary adaptation in new environments. Functional Ecology, 21(3):394–407, 2007. 10.1111/j.1365-2435.2007.01283.x.

Ghosh, D. D., Lee, D., Jin, X., Horvitz, H. R., & Nitabach, M. N. C. elegans discriminates colors to guide foraging. Science, 371(6533):1059–1063, 2021. 10.1126/science.abd3010.

Gillespie, J. Natural selection for within-generation variance in offspring number. Genetics, 76:601–606, 1974. 10.1093/genetics/81.2.403.

Guzella, T., Dey, S., Chelo, I. M., Pino-Querido, A., Pereira, V., Proulx, S., & Teotónio, H. Slower environmental change hinders adaptation from standing genetic variation. PLoS Genet, 14:e1007731, 2018. 10.1371/journal.pgen.1007731.

Heard, E. & Martienssen, R. A. Transgenerational epigenetic inheritance: myths and mechanisms. Cell, 157(1):95–109, 2014. 10.1016/j.cell.2014.02.045.

Hubbard, E. J., Korta, D. Z., & Dalfó, D. Physiological control of germline development. Adv Exp Med Biol, 757:101–31, 2013. 10.1007/978-1-4614-4015-45.

Jablonka, E., Oborny, B., Molnár, I., Kisdi, E., Hofbauer, J., & Czárán, T. The adaptive advantage of phenotypic memory in changing environments. Proceedings of the Royal Society B-Biological Sciences, 350:133–141, 1995. 10.1098/rstb.1995.0147.

Joschinski, J. & Bonte, D. Transgenerational plasticity and bet-hedging: A framework for reaction norm evolution. Frontiers in Ecology and Evolution, 8, 2020. 10.3389/fevo.2020.517183.

Kuijper, B. & Hoyle, R. B. When to rely on maternal effects and when on phenotypic plasticity? Evolution, 69:950–968, 2015. 10.1111/evo.12635.

Kuijper, B. & Johnstone, R. A. Evolution of epigenetic transmission when selection acts on fecundity versus viability. Philosophical Transactions of the Royal Society B: Biological Sciences, 376(1826):20200128, 2021. 10.1098/rstb.2020.0128.

Kuijper, B., Johnstone, R., & Townley, S. The evolution of multivariate maternal effects. PLoS Computational Biology, 10:e1003550, 2014. 10.1371/journal.pcbi.

LaMacchia, J., Frazier III, H., & Roth, M. Glycogenfuels survival during hyposmotic-anoxic stress in caenorhabditis elegans. Genetics, 201:65–74, 2015. 10.1534/genetics.115.179416.

Lande, R. Expected relative fitness and the adaptive topography of fluctuating selection. Evolution, 61(8):1835–46, 2007. 10.1111/j.1558-5646.2007.00170.x.

Lande, R. Adaptation to an extraordinary environment by evolution of phenotypic plasticity and genetic assimilation. J Evol Biol, 22(7):1435–46, 2009. 10.1111/j.1420-9101.2009.01754.x.

Lande, R. & Kirkpatrick, M. Selection response in traits with maternal inheritance. Genet Res, 55:189–197, 1990. 10.1017/s0016672300025520.

Lande, R. & Price, T. D. Genetic correlations and maternal effect offspringparent regression. Coefficients obtained from Genetics, 122:915–922, 1989.10.1093/genetics/122.4.915.

Leimar, O. & McNamara, J. The evolution of transgenerational integration of information in heterogeneous environments. Am Nat, 185:E55–E69, 2015. 10.1086/679575.

Liu, J., Ward, A., Gao, J., Dong, Y., Nishio, N., Inada, H., Kang, L., Yu, Y., Ma, D., Xu, T., Mori, I., Xie, Z., & Xu, X. Z. C. elegans phototransduction requires a g protein-dependent cgmp pathway and a taste receptor homolog. Nat Neurosci, 13(6): 715–22, 2010. 10.1038/nn.2540.

Lynch, M. & Walsh, B. Genetics and Analysis of Quantitative Traits. Sinauer Associates, Inc., Sunderland, 1998.

Marshall, D. & Uller, T. When is a maternal effect adaptive? Oikos, 116(12):1957–1963, 2007. 10.1111/j.2007.0030-1299.16203.x.

McElreath, R. Statistical Rethinking: A Bayesian Course with Examples in R and STAN. Chapman and Hall/CRC, 2nd edition, 2020. 10.1201/9780429029608.

McGlothlin, J. W. & Galloway, L. F. The contribution of maternal effects to selection response: an empirical test of competing models. Evolution, 68(2):549–58, 2014. 10.1111/evo.12235.

McNamara, J. M., Dall, S. R., Hammerstein, P., & Leimar, O. Detection vs. selection: integration of genetic, epigenetic and environmental cues in fluctuating environments. Ecol Lett, 19 (10):1267–76, 2016. 10.1111/ele.12663.

Mendenhall, A., LeBlanc, M., Mohan, D., & Padilla, P. A. Reduction in ovulation or male sex phenotype increases long-term anoxia survival in a daf-16-independent manner in caenorhabditis elegans. Physiol. Genomics, 36:167–178, 2009. 10.1152/physiolgenomics.90278.2008.-Identifying.

Michel, M. J., Chevin, L. M., & Knouft, J. H. Evolution of phenotype-environment associations by genetic responses to selection and phenotypic plasticity in a temporally autocorrelated environment. Evolution, 68(5):1374–84, 2014. 10.1111/evo.12371.

Moran, D. T., Dias, G. M., & Marshall, D. J. Associated costs and benefits of a defended phenotype across multiple environments. Functional Ecology, 24(6):1299–1305, 2010. 10.1111/j.1365-2435.2010.01741.x.

Moran, N. A. The evolutionary maintenance of alternative phenotypes. The American Naturalist, 139(5):971–989, 1992. 10.1086/285369.

Mousseau, T. & Fox, C. W. Maternal Effects as Adaptations. Oxford University Press, Oxford, 1998.

Nielsen, M. E. & Papaj, D. R. Why study plasticity in multiple traits? new hypotheses for how phenotypically plastic traits interact during development and selection. Evolution, 76(5): 858–869, 2022. 10.1111/evo.14464.

Noble, L. M., Chelo, I., Guzella, T., Afonso, B., Riccardi, D. D., Ammerman, P., Dayarian, A., Carvalho, S., Crist, A., Pino-Querido, A., Shraiman, B., Rockman, M. V., & Teotónio, H. Polygenicity and epistasis underlie fitness-proximal traits in the caenorhabditis elegans multiparental experimental evolution (cemee) panel. Genetics, 207(4): 1663–1685, 2017. 10.1534/genetics.117.300406.

Otto, S. P. & Day, T. A Biologist’s Guide to Mathematical Modeling in Ecology and Evolution. Princeton University Press, Princeton, 2007.

Pearl, J. Causality: Models, Reasoning, and Inference. Cambridge University Press, 2 edition, 2009. ISBN 9780521895606.

Proulx, S. & Day, T. What can invasion analyses tell us about evolution under stochasticity in finite populations? Selection, 2:1–15, 2001. 10.1556/Select.2.2001.1-2.2.

Proulx, S. & Teotónio, H. What kind of maternal effects are selected for in fluctuating environments? American Naturalist, 189(6):E118–E137, 2017. 10.1101/034546.

Proulx, S., Dey, S., Guzella, T., & Teotónio, H. How differing modes of non-genetic inheritance affect population viability in fluctuating environments. Ecology Letters, 22:1767–1775, 2019. 10.1111/ele.13355.

R Core Team. R: A Language and Environment for Statistical Computing. R Foundation for Statistical Computing, Vienna, Austria, 2021.

Roach, D. & Wulff, R. Maternal effects in plants. Ann Rev Ecol Evol Syst, 18:209–235, 1987. 10.1146/annurev.es.18.110187.001233.

Saether, B. E. & Engen, S. The concept of fitness in fluctuating environments. Trends Ecol Evol, 30(5):273–81, 2015. 10.1016/j.tree.2015.03.007.

Scheiner, S. M. The genetics of phenotypic plasticity. xvi. interactions among traits and the flow of information. Evolution, 72(11):2292–2307, 2018. 10.1111/evo.13601.

Seger, J. & Brockmann, H. J. What is bet-hedging? In Harvey, P. H. & Partridge, L., editors, Oxford Surveys in Evolutionary Biology, volume 4, pages 182–211. Oxford University Press, Oxford, 1987.

Simons, A. M. Modes of response to environmental change and the elusive empirical evidence for bet hedging. Proc Biol Sci, 278 (1712):1601–9, 2011. 10.1098/rspb.2011.0176.

Stan Development Team. RStan: the R interface to Stan, 2020. R package version 2.21.2.

Stewart, A. D. & Phillips, P. C. Selection and maintenance of androdioecy in caenorhabditis elegans. Genetics, 160(3):975–982, 2002. 10.1093/genetics/160.3.975.

Teotónio, H., Carvalho, S., Manoel, D., Roque, M., & Chelo, I. M. Evolution of outcrossing in experimental populations of Caenorhabditis elegans. PloS one, 7(4):e35811, 2012. 10.1371/journal.pone.0035811.

Teotónio, H., Estes, S., Phillips, P., & Baer, C. Experimental evolution with caernohabditis nematodes. Genetics, 206(12): 691–716, 2017. 10.1534/genetics.115.186288.

Theologidis, I., Chelo, I. M., Goy, C., & Teotónio, H. Reproductive assurance drives transitions to self-fertilization in experimental caenorhabditis elegans. BMC Biology, 12(1):93, 2014. 10.1186/s12915-014-0093-1.

Tufto, J. The evolution of plasticity and nonplastic spatial and temporal adaptations in the presence of imperfect environmental cues. The American Naturalist, 156(2):121–130, 2000. 10.1086/303381.

Tufto, J. Genetic evolution, plasticity, and bet-hedging as adaptive responses to temporally autocorrelated fluctuating selection: A quantitative genetic model. Evolution, 69(8):2034–49, 2015. 10.1111/evo.12716.

Via, S., Gomulkiewicz, R., De Jong, G., Scheiner, S. M.,Schlichting, C. D., & Van Tienderen, P. H. Adaptive phenotypic plasticity: consensus and controversy. Trends Ecol Evol, 10:212–217, 1995. 10.1016/S0169-5347(00)89061-8.

Wade, M. J. & Kalisz, S. The causes of natural selection. Evolution, 44(8):1947–1955, 1990. 10.1111/j.1558-5646.1990.tb04301.x.

Walsh, B. & Lynch, M. Evolution and Selection of Quantitative Traits. Sinauer Associates, Inc., Sunderland, 2018.

Ward, A., Liu, J., Feng, Z., & Xu, X. Z. Light-sensitive neurons and channels mediate phototaxis in c. elegans. Nat Neurosci, 11(8):916–22, 2008. 10.1038/nn.2155.

Wolf, J. B. Gene interactions from maternal effects. Evolution, 54(6):1882–98, 2000. 10.1111/j.0014-3820.2000.tb01235.x.

Wolf, J. B. & Brodie, E. D., r. The coadaptation of parental and offspring characters. Evolution, 52(2):299–308, 1998. 10.1111/j.1558-5646.1998.tb01632.x.

Wolf, J. B. & Wade, M. J. What are maternal effects (and what are they not)? Philos Trans R Soc Lond B Biol Sci, 364(1520): 1107–15, 2009. 10.1098/rstb.2008.0238.

Wolf, J. B. & Wade, M. J. Evolutionary genetics of maternal effects. Evolution, 70(4):827–39, 2016. 10.1111/evo.12905.

Wong, T. G. & Ackerly, D. D. Optimal reproductive allocation in annuals and an informational constraint on plasticity. New Phytologist, 166(1):159–172, 2005. 10.1111/j.1469-8137.2005.01375.x.

Zhang, W., He, F., Ronan, E. A., Liu, H., Gong, J., Liu, J., & Xu, X. S. Regulation of photosensation by hydrogen peroxide and antioxidants in c. elegans. PLOS Genetics, 16(12):1–21, 12 2020. 10.1371/journal.pgen.1009257.

Zhou, L. & Liu, Y. Light sensing enhances thermotolerance and competitive fitness via serotonergic signaling in an eyeless organism. Cell Research, 36(4):286–299, 2026. 10.1038/s41422-026-01223-x.

Zilio, G., Bedhomme, S., Fronhofer, E. A., Jacob, S., Legrand, D., Philippe, H., & Chevin, L.-M. Addressing multi-generational non-genetic responses in experimental studies of evolution. Evolution, page qpag027, 2026. 10.1093/evolut/qpag027.

