## supplementary materials for "Cue reliability and the evolution of adaptive plasticity in maternal traits"

### Supplementary Methods and Figures for: Cue reliability and the evolution of adaptive plasticity in maternal traits

### Detailed Materials and Methods

#### Ancestral population and standard culture conditions

All populations employed for experimental evolution are derived from a single ancestral population (named GA250). This ancestral population was derived from the hybridization of 16 inbred founder strains (Teotónio et al., 2012; Teotónio et al., 2017), followed by 140 generations of domestication to lab conditions (Teotónio et al., 2012; Chelo and Teotónio, 2013; Carvalho et al., 2014) and further 50 generations of gradual adaptation to high salt conditions, 305 mM of NaCl in growth media, from L1 larval stage to reproduction (Theologidis et al., 2014; Chelo et al., 2019).

Standard culture conditions during domestication and high salt evolution included maintaining populations at sizes of $N = 10^4$, and effective population sizes of $N_e = 10^3$ (Chelo and Teotónio, 2013), under a 4 day life-cycle (Teotónio et al., 2012). Each population was kept in ten Petri plates with solid NGM-lite agar media (Europe Bioproducts) covered by an overnight grown lawn of HT115 *E. coli* that served as *ad libitum* food. After growth to maturity for 66 hours at constant 20$^o$C and 80% relative humidity, nematodes from all 10 plates were harvested with M9 isotonic solution and exposed to 1 M KOH: 5% NaOCl bleach solution for 5 min, to which only embryos survive. These embryos were then kept in M9 without *E. coli* for 18 hours. After this period, live L1 stage larvae density was estimated (five 5 $\mu$L drops multiplied by total M9 volume), with the appropriate number of L1s then seeded to initiate the next generation. Under these culture conditions generations are non-overlapping.

*C. elegans* hermaphrodites are capable of self-fertilization but of outcrossing only when mated with males (Stewart and Phillips, 2002). During domestication, males were maintained at a frequency of 20%-40% and thus outcrossing rates were between 40%-80% (Teotónio et al., 2012; Carvalho et al., 2014). High salt greatly reduces mating chances such that the ancestral population was almost devoid of males after high salt adaptation (Theologidis et al., 2014). At the start of experimental evolution reported here, and despite predominant selfing, there was standing genetic variation for evolutionary responses to result from the sorting of selfing lineages (Noble et al., 2017; Dey et al., 2016). Genome-wide SNP diversity estimates of the number of effective lineages segregating in the ancestral population, as defined by the inverse of the sum of squared lineage frequencies, were of 30 (for total number of at least 250, see Guzella et al. (2018) for this analysis of the ancestral population). A subset of 4 unreliable populations (U3, U4, U7 and U8, see below) were previously reported and showed an effective number of three lineages segregating after the 60 generations of experimental evolution [see Figure S3 in Dey et al. (2016)].

#### Oxygen and blue light culture conditions

The first environmental factor manipulated during experimental evolution, and in the fitness and fecundity assays (see below), was atmospheric oxygen levels. Anoxic or normoxic conditions were imposed from early embryogenesis, after adult bleach, until the larvae L1 stage (Supplementary Figure S1). Embryonic DNA transcription, RNA translation, and other metabolism can start early, although it is known that maternal RNA, protein, and other metabolite contributions are required at least until gastrulation and embryonic survival in anoxia almost exclusively depends on maternal glycogen provisioning (Bowerman, 1998; Frazier and Roth, 2009; LaMacchia et al., 2015).

In detail, after bleaching and repeated washes with M9, 200 $\mu$L containing embryos were transferred to 25 mM NaCl NGM-lite plates, without *E. coli*. Petri plates were then placed inside 7 L polycarbonate boxes with rubber clamp-sealed lids (AnaeroPack, Mitsubishi Inc.). Within these boxes, anoxic conditions were imposed by placing two GasPak™ EZ sachets (Becton, Dickinson and Company, ref. no. 260678). Anoxic conditions, defined as $< 1\%$ $O_2$ or approximately 1 kPa $O_2$ at atmospheric pressure according to the manufacturer, were confirmed using BBL™ Dry Anaerobic Indicator Strips (Becton, Dickinson and Company, ref. no. 271051). To prevent desiccation, paper towels moistened with dd$H_2O$ were placed inside each box. For normoxic conditions, the GasPak™ EZ sachets were not used. After 16 hours, boxes were opened and live L1 larvae were washed off the plates with M9 buffer, and their density was estimated to proceed with the next generation.

The second environmental factor manipulated was light exposure during oogenesis and ovulation and prior to reproduction. Adult *C. elegans* are known to avoid light (Ward et al., 2008; Edwards et al., 2008), though in our setup, individuals could not escape the light source. We assume that all individuals received the same amount of light. Petri plate rack holders (Starsted) were fitted with strips of blue light LEDs (Nichia NS6B083T and NSPB300B), positioning each plate to receive illumination from approximately 10 individual LEDs placed on opposing sides. According to the manufacturer, these LEDs have a peak relative intensity at 465 nm. Accounting for Petri plate geometry, as well as reflection and scattering from plastic surfaces, the resulting irradiance was approximately 0.07 mW mm$^{-2}$ and 21,000 lux per plate. Starting 48 h after L1 seeding and over a 12 h period, blue light was flashed for 0.5 seconds every 2 seconds to prevent heat accumulation. A custom Python script running on a dedicated computer synchronized light exposure across all plates, while unexposed plates were kept in a separate incubator.

#### Experimental evolution design

*C. elegans* can be cryopreserved (Stiernagle, 1999), with stocks being frozen at $-80\,^{\circ}$C and revived as needed for experimental evolution and for comparison of ancestral and derived populations in common garden assays (Teotónio et al., 2017). The ancestral GA250 population was thawed from $-80\,^{\circ}$C stocks with $> 10^4$ individuals and cultured for two generations for expansion of individual numbers ($> 10^5$). GA250 was then divided into several populations to undergo experimental evolution (Supplementary Figure S2).

The sequences of fluctuating normoxia and anoxia during the 60 generations have been described in Proulx et al. (2019). Briefly, oxygen level sequences were designed so that there were 30 generations in normoxia and 30 generations in anoxia and a weak negative temporal autocorrelation of anoxic conditions between maternal and offspring generations ($\rho_1$=-0.065). We used three sequences so that the anoxia autocorrelation between grandmaternal and grand-offspring generations were close to zero ($\rho_2$=-0.017), positive ($\rho_2$=0.247) or negative ($\rho_2$=-0.311), and named Seq19, Seq31 and Seq33, respectively (Supplementary Figure S2). Temporal autocorrelations at higher lags were negligible.

For each oxygen sequence, hermaphroditic mothers either reliably or unreliably received blue light pulses depending on whether their progeny would face anoxia (Supplementary Figure S2). For the unreliable regime, the same number of 30 generations were subject to blue light pulses, although there was only 50% chance of cueing anoxia in the following generation. The temporal cross-correlation between light exposure in maternal generations and anoxia exposure in offspring generations was thus of one ($R_1$=1) in the reliable regime and of zero ($R_1$=0) in the unreliable regime, while keeping the same number of anoxia and light generations across the 60 generations in both regimes. There were weak cross-correlations between light in grandmaternal generations and anoxia in grand-offspring generations (Supplementary Figure S2): for unreliable regimes in Seq19, Seq31 and Seq33 they were of $R_2$=0.153, $R_2$=0.085, and $R_2$=-0.05, respectively; for reliable regimes they were of $R_2$=-0.085 for all oxygen sequences. Temporal cross-correlations at higher lags were negligible.

For each oxygen and light sequence of environmental states and reliability regime, we independently cultured 3 replicate populations during the 60 generations, following the standard laboratory conditions with high salt (305 mM NaCl) in the growth media and population sizes of N=$10^4$ from the L1 larval stage until reproduction. A total of 18 populations were maintained: for Seq19 there were replicate populations U3, U4 and U10 (in the 'Unreliable' light regime populations), and replicate populations R4-R6 (in the 'Reliable' light regime populations); for sequence Seq33, U16-U18 and R16-R18; and for sequence Seq31, U7, U8, U12, and R10-R12. A larger set of populations underwent experimental evolution but they were not assayed (see Proulx et al., 2019).

#### Fitness and fecundity assays

The discrete-time population growth multiplier, which we call $R_0$, was used as a proxy of fitness in anoxic conditions (see next section). It was measured by dividing the number of live L1-staged larvae after anoxia exposure, and after oxygen level manipulation in the grandmaternal and maternal generations, and light treatment in the maternal generation, by the fixed number of L1 seeded in the maternal generation (Supplementary Figure S1).

We measured $R_0$ in ancestral and evolved populations in common garden assays, after the experiment was finished. Assays were done in 18 different blocks, each corresponding to the date of thawing population samples and assay culture. In each block, all oxygen and light manipulations were done (8 conditions) for the ancestral GA250 population and one of the 18 evolved populations. All environmental conditions were the same as those during experimental evolution. Starting in the grandmaternal generation, three technical replicates for each population sample were run in parallel in each block.

In more detail, we thawed at least $10^3$ L1 individuals from $-80\,^{\circ}$C stocks to regular Petri plates (25 mM NaCl, with *E. coli*, normoxia) and surviving adults bleached to derive one culture per population sample. These were then maintained for one full generation alongside in standard conditions for individual number expansion. On the third generation, after bleach, each culture was split into three cultures (each with 5000 individuals, 5 Petri plates), with each culture being then in either normoxia or anoxia (starting the grandmaternal generation). Adults were bleached and individuals from the maternal generations were then exposed to varying oxygen during development and initial larval growth and varying light during early adulthood (5000 individuals, 5 Petri plates). After bleaching the maternal generation, embryos and L1 larvae from the offspring generation were exposed to anoxia. The total number of surviving offspring L1 was measured by counting them in ten 5 $\mu$L drops, scaled up to the total volume of the wash (2-3 mL, to the nearest $\mu$L). This number was then divided by 5000 to obtain the population sample $R_0$ in anoxia (see next section on fitness analysis and Supplementary Figure S1).

A random id was assigned to each technical replicate to diminish potential bias during handling. All assays were performed by the same experimenter. There were therefore 5 generations of common culture between samples, which precludes our estimates from being biased due to uncontrolled environmental effects up to 5 generations. Due to unavoidable adult (prior to bleaching) and live L1 (after bleaching) loss that occurred with our manipulations, the reported $R_0$ are a lower bound measure of the true values. Overall, the data for analysis consists of 845 observations: 18 blocks each with one evolved population plus the ancestral population, times 4 oxygen conditions, times 2 light conditions, times 3 technical replicates, minus accidental loss because of insufficient L1 numbers for continued culture in grandmaternal or maternal generations.

We ran fecundity assays in 6 thaw-date blocks, separately from the fitness assays, each including the ancestor population and one evolved population from reliable or unreliable regimes of Seq31 (R10, R11, R12; U7, U8, U12). Fecundity was measured as the number of embryos after grandmaternal and maternal oxygen and light manipulations, and thus before progeny exposure to anoxia as in the fitness assays (Supplementary Figure S1), divided by the fixed number of seeded L1 larvae in the maternal generation. All other design details were similar to the fitness assays (technical replication, id randomization, etc.). We thus assume throughout that all populations have the same individual viability from larvae to reproduction. The data for analysis consists of 285 observations. Reported fecundity estimates are a lower bound measure of the true values because of adult and embryo loss prior to counting.

#### Fitness analysis of the ancestral population

The discrete-time population growth multiplier, $R_0$, is surrogate for fitness that was measured in our non-overlapping maternal-offspring generation experimental evolution protocol, with the natural log of the $R_0$ being roughly equivalent to the instantaneous population growth rate (Otto and Day, 2007). Given the experimental evolution regimes, changes in $w = \ln R_0$ should represent changes in Malthusian fitness per generation, and we conduct the statistical analyses on this measure. Note, however, that for the figures we back-transformed $w$ to show the results in the $R_0$ scale.

We first defined an index variable for the measurement block of each assay observation, where $b[i]$ defines the block in which the $i$th observation was made. Next, we defined an index variable for the assay state of each experimental observation, where

$s[i]$ defines the state of the $i$th observation in our data table. We uniquely defined the states with an index from 1 through 8 to represent all possible combinations of maternal normoxia/anoxia, grandmaternal normoxia/anoxia, and maternal light/no-light exposure.

We first fit the ancestral population to better understand the baseline the experimental populations were starting from, using the following Bayesian model (results shown in Figure 1):

$$\begin{aligned} w_i &\sim \mathcal{N}(\mu_i,\, \sigma_{\text{obs}}) \\ \mu_i &= \alpha_A + \eta_{A,\, s[i]} + \epsilon_{\text{block},\, b[i]} \\ \epsilon_{\text{block},\, b} &\sim \mathcal{N}(0,\, \sigma_{\text{block}}) \end{aligned} \tag{S1}$$

where $i$ refers to the observation number, $w_i$ is (Malthusian) fitness, $\mu_i$ is the expected fitness, $\alpha_A$ is the overall mean fitness of the Ancestral population, and $\eta_{A,s[i]}$ is the effect of state $s[i]$ on ancestral fitness. The term $\epsilon_{\text{block},\, b[i]}$ represents a random effect of assay block which is normally distributed with standard deviation $\sigma_{\text{block}}$, and $\sigma_{\text{obs}}$ is the standard deviation of the normal likelihood function. Estimating $\eta_{A,s[i]}$ independently for each state is equivalent to including all interactions among maternal oxygen environment, grandmaternal oxygen environment, and maternal light environment.

In addition to the likelihood and hierarchical prior structure of the model, we imposed a sum-to-zero constraint on the state effects $\eta_{A,s}$, such that $\sum_s \eta_{A,s} = 0$. Centering the random effects $\epsilon$ at zero ensures that they do not shift the overall mean, making the intercept $\alpha_A$ directly interpretable as the ancestral population mean fitness. Without a constraint on $\eta_{A,s}$, the model would be non-identifiable, as adding a constant to $\alpha_A$ and subtracting it from all of the state effects would not change the likelihood (but would alter the calculated hyper prior and priors).

In this model, each observation of fitness is assumed to follow a normal distribution which can be thought of as encompassing both random variation in birth rates and sampling variance. The measurement block is also given a random effect, where the variance among blocks is determined by an adaptive prior on the variance between blocks. From this model we can estimate fitness under all 8 environmental states along with the uncertainty in our estimate (McElreath, 2020).

#### Fitness analysis of reliable and unreliable populations

We next fit a model to infer the evolved fitness changes from the ancestral population, which we call evolutionary *divergence*, and partition these changes into effects related to experimental evolution regime and environmental state. We represent evolved populations as deviations from the ancestral mean fitness and ancestral state-specific effects. This formulation allows us to separately estimate evolutionary changes in overall mean fitness and in the environmental state-dependent fitness responses that are forms of fitness plasticity for the oxygen and light assay environments. We term these differences between the reliable and unreliable populations, relative to the ancestral population, as evolutionary *differentiation*.

Indicator variables were defined for the evolutionary regime, $I_{R,i}$ and $I_{U,i}$ which take a value of 1 if observation $i$ is from a reliable or unreliable evolved population, respectively, and 0 otherwise. Another indicator variable $I_{E,i}$ takes a value of 1 for evolved populations (Reliable or Unreliable) and 0 for ancestral populations. Likewise, index variables for replicate population and the evolutionary sequence of environments were defined (Seq19, Seq31, Seq33, see above), which only apply to evolved populations, where $j[i]$ indexes replicate population and $q[i]$ indexes the sequence number. Note that population index is unique for all evolved populations, but this index does not relate to which regime the population is from. However, when sum-to-zero constraints are enforced they are applied separately for each experimental evolution regime.

The model builds on the ancestral model (equation (S1)) with all of the common terms using the same definition and is:

$$
\begin{aligned}
w_i &\sim \mathcal{N}(\mu_i,\, \sigma_{\text{obs}}) \\
\mu_i &= \alpha_A + \eta_{A,\,s[i]} + \Delta\alpha_R\, I_{R,i} + \Delta\alpha_U\, I_{U,i} \\
&\quad + \Delta\eta_{R,\,s[i]}\, I_{R,i} + \Delta\eta_{U,\,s[i]}\, I_{U,i} \\
&\quad + \epsilon_{\text{block},\,b[i]} + \epsilon_{\text{seq},\,q[i]}\, I_{E,i} + \epsilon_{\text{pop},\,j[i]}\, I_{E,i} \\
\epsilon_{\text{block},\,b} &\sim \mathcal{N}(0,\, \sigma_{\text{block}}) \\
\epsilon_{\text{seq},\,q} &\sim \mathcal{N}(0,\, \sigma_{\text{seq}}) \\
\epsilon_{\text{pop},\,j} &\sim \mathcal{N}(0,\, \sigma_{\text{pop}})
\end{aligned}
\tag{S2}
$$

In this model, $\Delta\alpha_R$ and $\Delta\alpha_U$ represent changes in fitness of the Reliable and Unreliable evolved populations relative to the ancestor. These terms represent the baseline changes in overall fitness irrespective of the environment due to evolution and can be interpreted as divergence from the ancestral population in the average "elevations" of the reaction norms in oxygen and light assay environments. The $\Delta\eta$ terms represent average evolutionary changes in the state-specific fitness for each reliability regime (results shown in Figure 2), and it will be contrasts among these that allow us to infer changes in the reaction norm slope for fitness particularly with respect to the light environment (see section below on contrasts).

The term $\epsilon_{\text{seq},\,q[i]}$ represents the effect of environmental state sequence during experimental evolution and only applies to evolved populations. This is an additive term and so each sequence has the same effect on reliable and unreliable population means. Sum-to-zero conditions are enforced. The term $\epsilon_{\text{pop},\,j[i]}$ represents the effect of replicate population, and only applies to evolved populations. Sum-to-zero conditions are enforced separately for reliable and unreliable populations, but a common $\sigma$ term is used. Thus variability among replicate populations is assumed to be statistically the same for reliable and unreliable populations, but separate enforcement of sum-to-zero conditions ensures that the $\Delta$ terms can be interpreted without accounting for the $\epsilon$ values.

As in the ancestral model, estimating $\eta_{A,s[i]}$, $\Delta\eta_{R,s[i]}$, and $\Delta\eta_{U,s[i]}$ independently for each state is equivalent to including all interactions among maternal oxygen environment, grandmaternal oxygen environment, and maternal light environment, as well as their evolutionary change. Put another way this model allows for interactions between maternal oxygen environment, grandmaternal oxygen environment, light environment, and experimental evolution regime. As in the ancestral model, fitness observations are assumed to follow a normal distribution, capturing both stochastic variation in birth and measurement error. Variation among assay blocks, evolutionary sequences, and replicate populations is modeled using hierarchical random effects with variances estimated in a partial-pooling fashion from the data.

Weakly informative but regularizing priors were used:

$$\begin{aligned}
\alpha_A &\sim \mathcal{N}(0,\, 1.5)\\
\Delta\alpha_R,\, \Delta\alpha_U &\sim \mathcal{N}(0,\, 1)\\
\eta_{A,s} &\sim \mathcal{N}(0,\, \sigma_{\eta_A})\\
\Delta\eta_{R,s},\, \Delta\eta_{U,s} &\sim \mathcal{N}(0,\, \sigma_{\Delta\eta})\\
\sigma_{\eta_A},\, \sigma_{\Delta\eta},\, \sigma_{\text{block}},\, \sigma_{\text{seq}},\, \sigma_{\text{pop}},\, \sigma_{\text{obs}} &\sim \text{Exp}(1).
\end{aligned}$$

Implementation included using a non-centered parameterization, in which standardized parameters were scaled by their corresponding standard deviation parameters prior to applying the sum-to-zero constraint. The model was fit using Hamiltonian Monte Carlo with six independent chains with 2,000 warm-up iterations followed by 4,000 sampling iterations (each). Convergence was assessed by verifying that all $\hat{R} < 1.01$, and bulk effective sample sizes exceeded 2,500 for all model parameters.

#### Fecundity analysis

Fecundity was modeled in a similar way to fitness, with two key differences. First, because only a single evolutionary environmental sequence was assayed (Seq31, see above), no sequence-level random effect was included. Second, we modeled fecundity using a Gamma likelihood with a log link function, which is a compromise between modeling fecundity as a fully discrete variable using a negative-binomial distribution and log transforming fecundity and using a normal likelihood. Each of these approaches has theoretical benefits depending on how variance scales with the mean values and whether or not the number of counted embryos is large. Instead of relying solely on theory for when these distributions converge on each other, we also ran a log-transformed normal likelihood model, and the fecundity estimates for each population sampled were almost identical (the average deviation was on the order of 0.1% and the variance in deviation across all observations was 0.001). The Gamma distribution is parameterized in terms of shape and rate, such that the expectation is given by the ratio of the shape to the rate.

We modeled the expected fecundity on the log scale as an additive function of the maternal/grandmaternal oxygen environment and the maternal light environment (presence/absence). This additive structure was used because we found no evidence for an interaction between oxygen and light environments in preliminary analyses. As in the fitness model, we used sum-to-zero constraints and hierarchical priors to ensure identifiability and regularization. We modeled the effects of oxygen environments and light environment separately for the ancestral, reliable, and unreliable populations so that we could separately test for effects and interactions within each experimental evolution regime. We coded light as a contrast-coded variable taking on the value of -1 for no light (dark) and 1 for light exposure.

The model specification is:

$$
\begin{aligned}
f_i &\sim \text{Gamma}(\phi,\ \phi/\mu_i)\,, \\
\log(\mu_i) &= \alpha_A I_{A,i} + \alpha_R I_{R,i} + \alpha_U I_{U,i} \\
&\quad + \eta_{A,M,\,m[i]} I_{A,i} + \eta_{R,M,\,m[i]} I_{R,i} + \eta_{U,M,\,m[i]} I_{U,i} \\
&\quad + \eta_{A,C} C_i I_{A,i} + \eta_{R,C} C_i I_{R,i} + \eta_{U,C} C_i I_{U,i} \\
&\quad + \epsilon_{\text{block},\,b[i]} + \epsilon_{\text{pop},\,j[i]} I_{E,i} \\
\epsilon_{\text{block},\,b} &\sim \mathcal{N}(0,\ \sigma_{\text{block}}) \\
\epsilon_{\text{pop},\,j} &\sim \mathcal{N}(0,\ \sigma_{\text{pop}}).
\end{aligned}
\tag{S3}
$$

Again, $I_{A,i}$, $I_{R,i}$, and $I_{U,i}$ are indicator variables denoting whether observation $i$ belongs to the Ancestral, Reliable, or Unreliable regime, and $I_{E,i}$ is an indicator variable for experimentally evolved populations. The parameters $\alpha_A$, $\alpha_R$, and $\alpha_U$ represent the log-scale fecundity intercept for each regime. The $\eta$ terms describe the effect of grandmaternal and maternal oxygen environments ($M$) or maternal light environment ($C$) and are set separately for each evolutionary regime including the ancestor. For oxygen environments, they also depend on which state $m[i]$ the observation was made in.

The $\epsilon$ terms represent variability in the hierarchical model, where we enforced the sum-to-zero conditions. For fecundity data we have a block effect and a replicate population effect (only for evolved populations). The population effect for reliable and unreliable populations shared a common standard deviation $\sigma_{\text{pop}}$, but had the sum-to-zero constraint imposed separately for each evolutionary treatment. This ensures that the $\alpha$ parameters measure the group mean parameter (i.e. this prevents covariance between parameters from masking a group effect). Finally, $\phi$ is the Gamma shape parameter governing variation around the mean fecundity.

We used broad but regularizing priors of:

$$
\begin{aligned}
\alpha_A,\ \alpha_R,\ \alpha_U &\sim \text{Normal}(0,1), \\
\eta_{A,M,m},\ \eta_{R,M,m},\ \eta_{U,M,m} &\sim \text{Normal}(0,0.3), \\
\eta_{A,C},\ \eta_{R,C},\ \eta_{U,C} &\sim \text{Normal}(0,0.3), \\
\sigma_{\text{block}},\ \sigma_{\text{pop}} &\sim Exp(1) \\
\phi &\sim \text{LogNormal}(1,0.3).
\end{aligned}
$$

Thus, the fitted fecundity model estimated separate intercepts, maternal-environment effects, and light environmental effects for the ancestral population, and for the reliable and unreliable experimental evolution regimes, while accounting for assay block-to-block variation and replicate variation among evolved populations (main results shown in Figure 4).

#### Contrasting evolutionary responses

The evolution of the mean elevations of the fitness reaction norms are modeled by the $\Delta\alpha_R$ and $\Delta\alpha_U$ terms in equation (S2) (the portion of the response that is environment independent) and by the $\Delta\eta_R$ and $\Delta\eta_U$ terms, also in equation (S2) (which are specific to each environmental state) (shown in Figure 2). To model the evolution of the fitness reaction norm slopes in the light environment, which we term *light-plasticity*, we contrasted the $\Delta\eta$ terms.

For each posterior draw, we computed predicted outcomes on the natural scale and defined the maternal fitness responses to light assay treatment within an experimental regime as the difference between light and no-light conditions. This is done by first computing the fitted linear predictor (equation (S2), shown in Figure 2):

$$\mu_{T,s} = \alpha_A + \eta_{A,s} + \Delta\alpha_T + \Delta\eta_{T,s},$$

where $T$ can be either Reliable or Unreliable, and $\Delta\alpha_T + \Delta\eta_{T,s}$ represents the total evolutionary divergence in fitness of the experimental evolution treatment (Reliable or Unreliable; Ancestral being zero) relative to the ancestral population for each oxygen and light environmental state, and with block effects, sequence effects, and replicate population effects all set to zero (their average value). The difference in fitness due to treatment is the sum of the change in the intercept term $\alpha$ plus the change in the state-specific response term, $\eta$.

For each posterior sample, we computed the light response for each oxygen environmental state subtracting the mean fitness without light-exposure from the mean fitness with light-exposure:

$$\Delta_{T,m} = \exp\bigl(\mu_{T,\, s(m,\text{Light})}\bigr) - \exp\bigl(\mu_{T,\, s(m,\text{No Light})}\bigr)\,. \tag{S4}$$

The total evolutionary change between the reliable and unreliable experimental regimes in terms of light-plasticity, relative to the ancestral, is then estimated as the difference between reliable and unreliable regimes for each oxygen environmental state and as the difference between regimes in the average over the 4 grandmaternal and maternal oxygen environmental states (results shown in Figure 3):

$$\Delta_{\text{overall}} = \frac{1}{4}\sum_{m=1}^{4}\left(\Delta_{R,m} - \Delta_{U,m}\right), \tag{S5}$$

computed for each sample from the posterior to create the posterior distribution of the evolutionary change in light-responses. This quantity represents a difference of differences on the natural scale, so a value of zero indicates no differentiation in light-plasticity between experimental evolution regimes.

For the fecundity contrasts, we used an additive model of grandmaternal and maternal oxygen environments and light environment, so we could consider the light-plasticity at the mean grandmaternal and maternal oxygen effects. For each posterior draw we compute:

$$\Delta_{\text{light}} = (\exp(\alpha_R + \eta_{R,C}) - \exp(\alpha_R - \eta_{R,C})) - (\exp(\alpha_U + \eta_{U,C}) - \exp(\alpha_U - \eta_{U,C}))\,, \tag{S6}$$

and then summarize the distribution over all samples in the posterior. This quantity is a difference of a difference, and so values of zero indicate that the response to light is the same for reliable and unreliable populations.

### Supplementary Figures

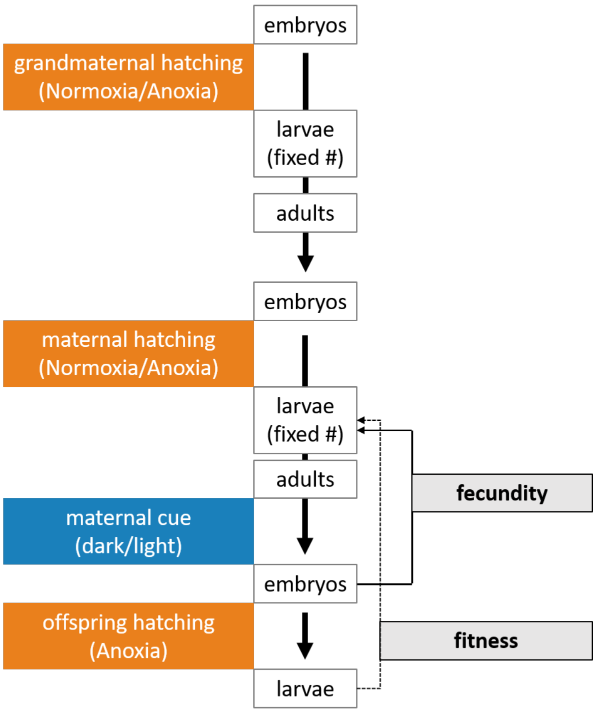


**Figure S1**: **Fitness and fecundity culture design.** Colored boxes indicate the environmental factors manipulated to assess their effects on maternal fecundity and offspring fitness in anoxia. Middle boxes represent the life-history stage during the grandmaternal, maternal, and early offspring generations. In the grandmaternal and maternal generations, embryos were exposed to either normoxic or anoxic hatching conditions (orange); in the offspring generation, only anoxic conditions were applied. Although we refer to anoxia as the absence of oxygen, we could only confirm that the oxygen concentration was below 1% in the atmosphere (see Materials and Methods). In the maternal generation, and prior to reproduction, adults were either exposed to blue light pulses or left in the dark (blue). The *per capita* population growth multiplier is calculated as the number of live hatched larvae following embryogenesis in anoxia, divided by the fixed number of larvae that initiated the maternal generation ($R_0$), which we consider to be a proxy for fitness in anoxia. Similarly, fecundity is measured as the number of embryos per capita before anoxic conditions in the offspring generation divided by the fixed number of larvae that initiated the maternal generation. For fecundity, we thus assume that all populations have the same individual viability from larvae to reproduction. All populations of experimental evolution, and the ancestral population from which they were derived, were measured contemporaneously in several assay blocks after 2 generations of common garden.

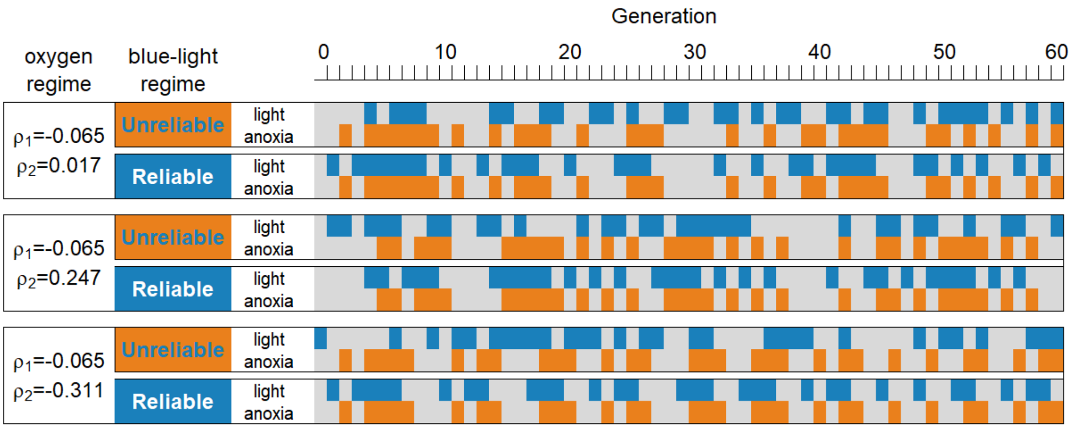


**Figure S2**: **Experimental evolution design.** Three environmental sequences with weak temporal autocorrelation of oxygen level conditions between maternal and offspring generations ($\rho_1$=-0.065; orange), and variable autocorrelations between grandmaternal and offspring generations ($-0.311 < \rho_2 < 0.247$), were imposed over 60 discrete non-overlapping generations. Maternal generations were exposed to blue light pulses that either reliably ($R_1$=1) or unreliably ($R_1$=0) cued anoxia in the offspring generation (blue). There were weak cross-correlations between grandmaternal light and offspring anoxia ($-0.08 < R_2 < 0.15$; see Materials and Methods). For each combination of oxygen and light environmental state sequences during experimental evolution, we cultured three independent replicate populations, for a total of 18 populations. All populations were derived at generation -2 from a single ancestral population with standing genetic variation, previously domesticated to standard culture conditions for 140 generations [$N = 10^4$, normoxia, no blue light pulses], followed by 50 generations of gradual adaptation to high salt (305 mM NaCl) in the larval-to-adult nematode-growth medium. Under high-salt conditions hermaphrodites are favored, and selfing was the predominant mode of reproduction during experimental evolution (see Materials and Methods for further details).

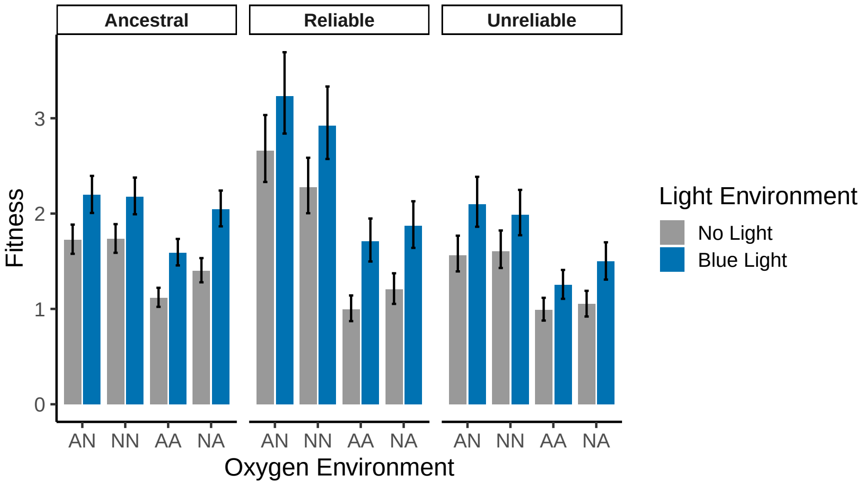


**Figure S3**: **Fitness of experimental populations.** Fitness under anoxic conditions was measured as the per capita population growth rate, $R_0$, following grandmaternal and maternal exposure to normoxia (N) or anoxia (A), and maternal exposure to either blue light pulses (blue) or no light (gray) during oogenesis and ovulation (Supplementary Figure S1). Measurements were made in 18 independent assay blocks, each with 3 technical replicates (Materials and Methods). The bars show posterior medians with 95% probable interval (PI) for the ancestral population (left), reliable populations (middle) and unreliable populations (right). Estimates should be interpreted as lower bounds on the true values, as nematode loss occurs during assay manipulations before counting (see Materials and Methods).

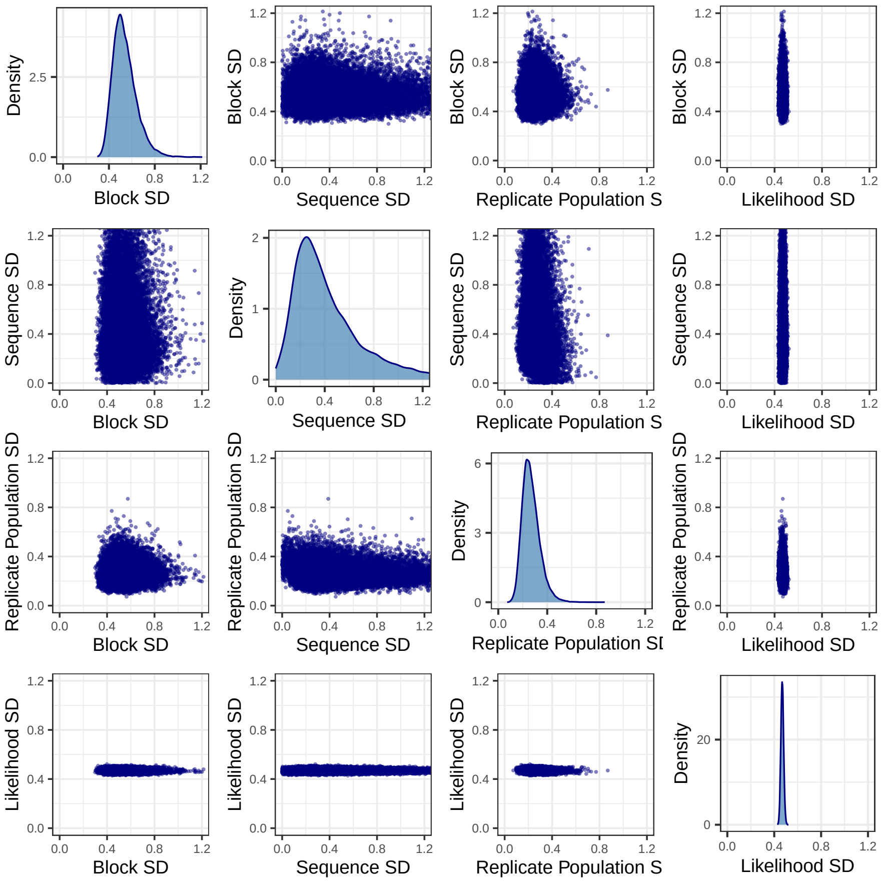


**Figure S4**: **Marginal and pair posterior distributions of variance terms in the fitness model.** The multilevel model involved three sources of variability (often termed random effects): assay statistical block, sequence of fluctuating normoxia-anoxia during experimental evolution, and replicate populations. Generically, these variance terms can trade off with each other to produce parameter sets with high likelihood. Each of the terms has a similar magnitude, although we have strong evidence that between-block variance is away from zero. The pair plots show, as expected, a negative correlation between each of the pairs of variance terms. The likelihood standard deviation is also shown.

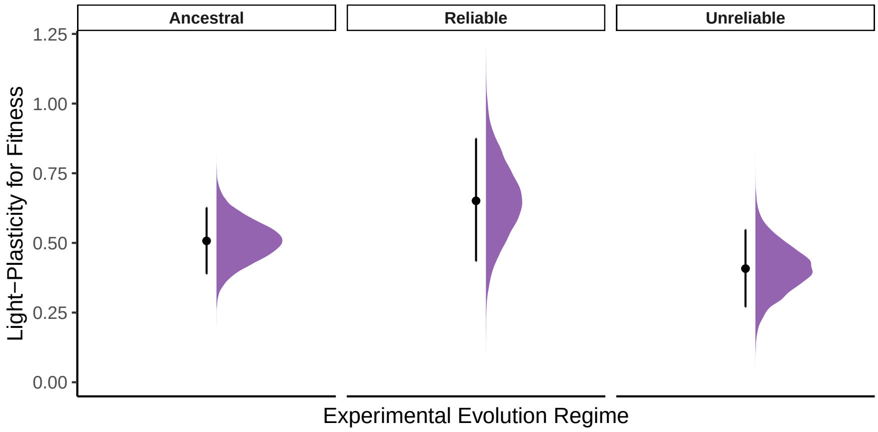


**Figure S5**: **Light-plasticity in fitness.** Light-plasticity is shown for each experimental evolution regime, averaged across grandmaternal and maternal oxygen environments (see equation (S5) in Materials and Methods). Shown are posterior medians with 95% probable intervals (PIs) for the ancestral population (left), reliable populations (middle) and unreliable populations (right). Half-eye plots show the posterior distribution estimates.

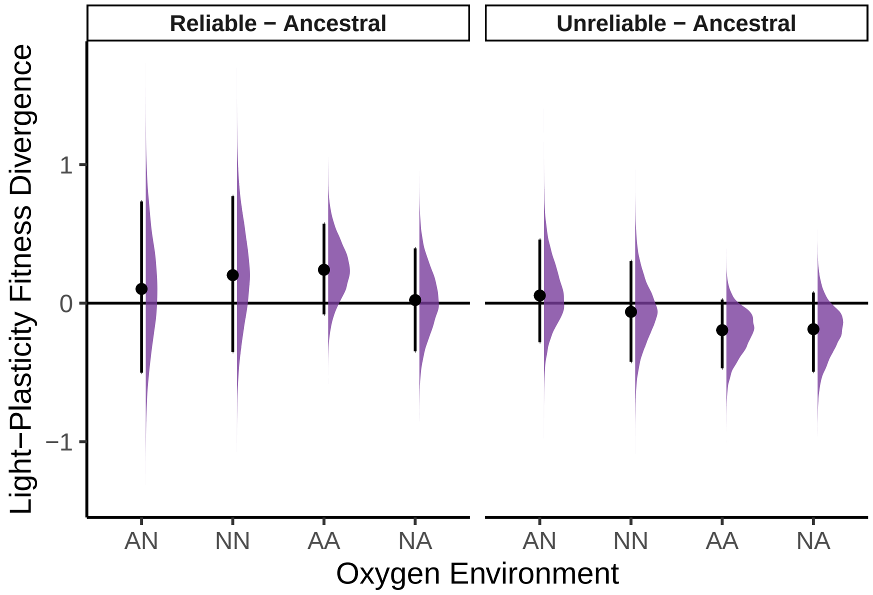


**Figure S6**: **Fitness divergence of light-plasticity.** Fitness difference between presence and absence of light of reliable (left) and unreliable (right) populations, relative to the ancestor population (see equation (S4) in Materials and Methods). Shown are posterior medians with the 95% probable interval (PI) with half-eye plots for the posterior distribution estimates. "N", normoxia; "A", anoxia in grandmaternal and maternal generations.

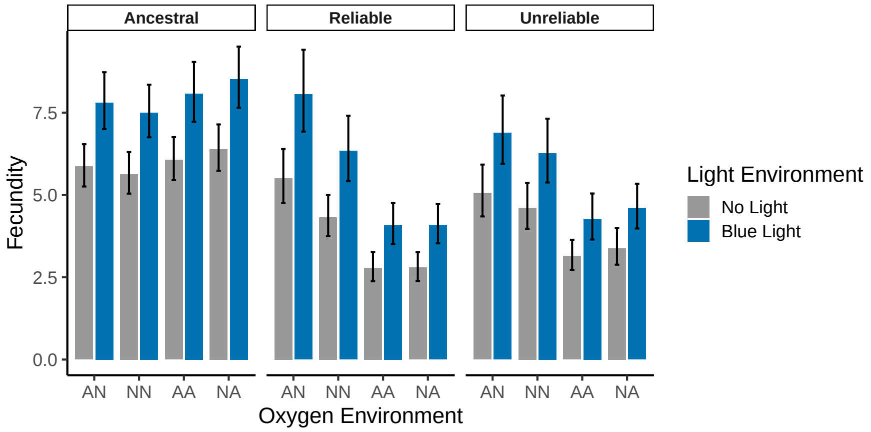


**Figure S7**: **Fecundity of experimental populations.** Fecundity was measured as the per capita embryo number, following grandmaternal and maternal exposure to normoxia (N) or anoxia (A), and maternal exposure to either blue light pulses (blue) or no light (gray) during oogenesis and ovulation (Supplementary Figure S1). Measurements were made in 6 independent assay blocks, each with 3 technical replicates (Materials and Methods). The bars show posterior medians with the 95% probable interval (PI) accounting for the uncertainty due to environmental state and light treatment, for the ancestral population (left), reliable populations (middle) and unreliable populations (right). Additional variability due to assay block and replicate population sum to zero and therefore do not contribute to uncertainty in the estimate. There however is additional uncertainty in the overall average fecundity (Supplementary Figure S9, Materials and Methods). Estimates should be interpreted as lower bounds on the true values, as nematode loss occurs during assay manipulations before counting.

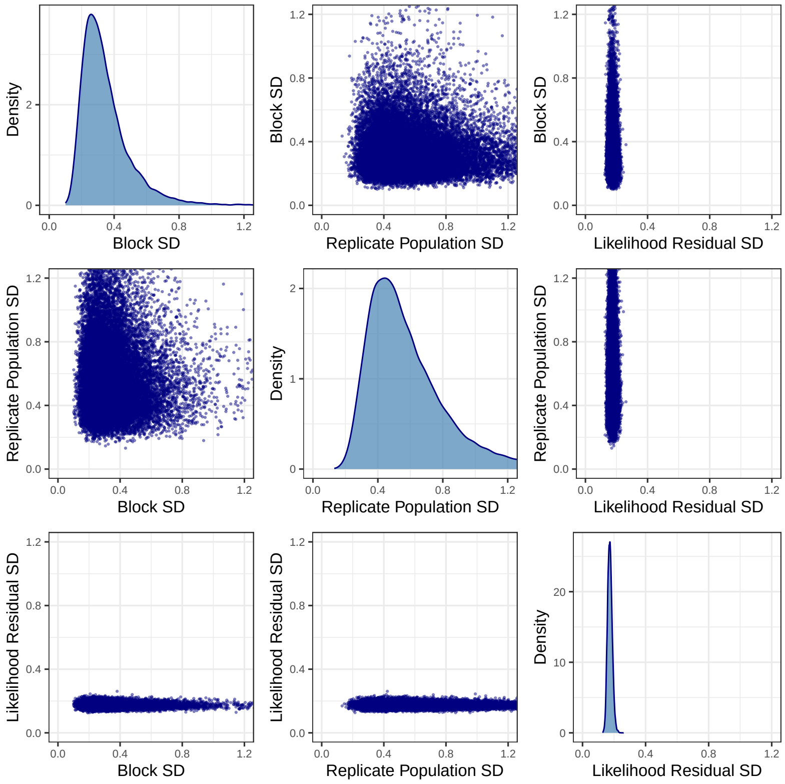


**Figure S8**: **Marginal and pair posterior distributions of variance terms in the fecundity model.** The multilevel model involved two sources of process variability: assay statistical block, and replicate populations. The likelihood standard deviation is also shown.

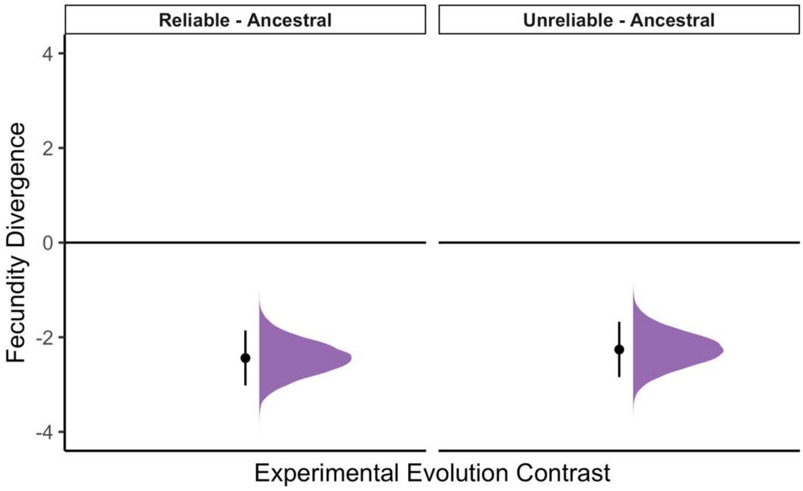


**Figure S9**: **Average fecundity regime divergence from ancestral.** Average fecundity divergence of reliable and unreliable populations from the ancestral. Shown is the posterior median with the 95% probable interval (PI), with the half-eye for the posterior distribution estimate. The contrast was calculated by taking the difference of the intercept fecundity parameter for each experimental regime, noting that all other effects are constrained to sum to zero across the maternal and light environments (Materials and Methods).

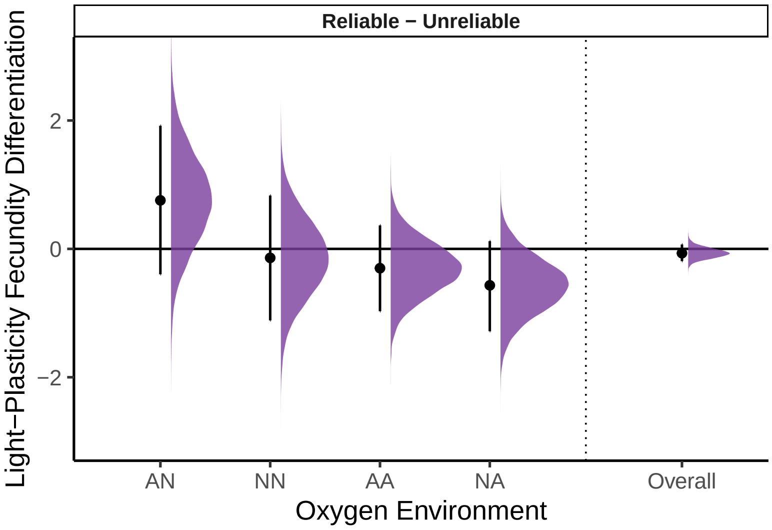


**Figure S10**: **Experimental regime differentiation of fecundity plasticity to the light environment.** Evolution of the fecundity difference between reliable and unreliable populations in light plasticity (presence versus absence), relative to the ancestral populations, with median and 95% PIs posterior estimates (dot and interval). Half-eye plots show the posterior distributions for each contrast (see equation (S6), Materials and Methods). Contrasts were calculated with the light plasticity set to 0, creating a slight bias due to the non-linear log link function. Including light effects would not change the sign of the contrasts and only slightly stretch their effects. "N" for normoxia, "A" for anoxia in grand-maternal and maternal generations.

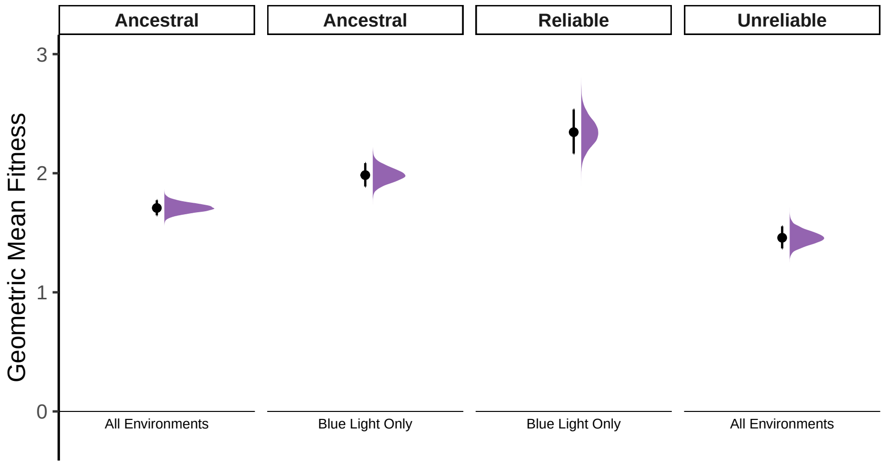


**Figure S11**: **Adaptation of experimental populations.** Geometric mean fitness estimates over all combinations of grandmaternal and maternal environments for reliable and unreliable populations (two right panels), calculated for the specific sequence of environments they were exposed to during experimental evolution. The ancestral population (two left distributions) shows the geometric mean fitness estimates had they had the same evolutionary history of environmental sequence states as the reliable (blue light only) or unreliable (all environments) populations. Shown is the posterior median with 95% probable interval (PI), with the half-eye for the posterior distribution estimate.
